# Seeing sounds: Neural mechanisms underlying auditory contributions to visual detection

**DOI:** 10.1101/2022.01.19.476187

**Authors:** Alexis Pérez-Bellido, Eelke Spaak, Floris P. de Lange

## Abstract

Sounds enhance the detection of visual stimuli while concurrently biasing an observer’s decisions. To investigate the neural mechanisms that underlie such multisensory interactions, we decoded time-resolved signal detection theory (SDT) sensitivity and criterion parameters from neural activity using magnetoencalography, while participants performed a visual detection task. Using temporal generalization analysis, we found that sounds improve visual detection by enhancing the maintenance of the most informative perceptual samples over time. In parallel, criterion decoding analyses revealed that sounds evoke patterns of activity that resembled the patterns evoked by an actual visual stimulus. These two complementary mechanisms of audiovisual interaction differed in terms of their automaticity: Whereas the sound-induced enhancement in visual information maintenance depended on participants being actively engaged in a detection task, sounds evoked visual activity patterns in the visual cortex in a bottom-up fashion, challenging the classical assumption that sound-induced reductions in criterion correspond to decision-level biases.

## Introduction

Although humans largely rely on vision in order to monitor the environment (Colavita, 1974; Posner et al., 1976), our brain has learnt that sensory signals from external events are often correlated across sensory modalities and should be optimally integrated (Ernst & Banks, 2002; Rohe & Noppeney, 2015). Conveniently, the human brain has adapted to a multisensory environment and in the last years multiple studies have revealed that regions classically considered as visual areas are tuned to sensory information conveyed by both visual and auditory signals (Deneux et al., 2019; Garner & Keller, 2021; Ibrahim et al., 2016; Murray et al., 2016). These results suggest that the processing of visual information might be already modulated by auditory inputs at the earliest perceptual stages (Driver & Noesselt, 2008). In visual detection, previous behavioral research has shown that synchronizing a sound with a visual stimulus improves the detection threshold of the latter. This phenomenon, termed as the sound induced visual enhancement, has been described using the Signal Detection Theory framework (Green & Swets, 1966) in multiple psychophysical studies (Fiebelkorn, Foxe, Butler, & Molholm, 2011; Frassinetti et al., 2002; Lippert et al., 2007). While task-irrelevant sounds improve participants visual sensitivity (d’) to discriminate signal from noise, they also increase the proportion of reported false alarms, leading to a reduction in the criterion parameter (c) (Frassinetti et al., 2002; Lippert et al., 2007; Lovelace et al., 2003; Odgaard et al., 2003; Pérez-Bellido et al., 2013). However, although the behavioral consequences of sounds on visual detection have been well characterized, we still lack a good description which specific neural mechanisms underly such multisensory perceptual decision-making biases in detection.

Most of previous neuroimaging research has focused on understanding how sounds enhance visual sensitivity. Nowadays, it is widely accepted that sounds modulate the processing of visual signals at early perceptual stages through sensory level interaction of audiovisual inputs (Giard & Peronnet, 1999; Mercier & Cappe, 2020; Meredith & Stein, 1983; Murray et al., 2016; van Atteveldt et al., 2014a). Yet, more recent studies tackling similar questions from a perceptual decision-making perspective (Franzen et al., 2020; Kayser et al., 2017) have characterized the effect of sounds on visual discrimination as a late enhancement in the transformation of sensory input into accumulated decisional evidence. Therefore, how specifically and to which extent sounds enhance visual sensitivity at early (i.e. sensory), late (i.e. decision) or both (Mercier & Cappe, 2020) perceptual stages remains under debate.

An equally relevant question that has received much less attention in perceptual neuroscience is how sounds induce changes in visual detection criterion. Although SDT criterion variations have been typically associated to decision level response biases, some perceptual illusions that affect perceptual accuracy often manifest in a shifted criterion parameter (J. Witt et al., 2012; J. K. Witt et al., 2015). Thus, whether the sound-induced criterion reduction reported in visual detection tasks exclusively reflects a decisional-level bias or it also depends on a perceptual-level bias is still unknown.

Finally, another disputed question taps into the automaticity of multisensory integration (Lunn et al., 2019; Macaluso et al., 2016; Talsma et al., 2010). While there is ample evidence that sounds can modulate neural activity in early visual areas in a bottom-up fashion, and these modulations may affect the processing of subsequent visual information (De Meo et al., 2015; Deneux et al., 2019; Ibrahim et al., 2016; Romei et al., 2009a, 2012), several studies have shown that crossmodal interactions are weakened when attention is directed away from the relevant stimuli (Alsius et al., 2007; Alsius & Soto-Faraco, 2011; Convento et al., 2018; Talsma et al., 2007; Talsma & Woldorff, 2005), or the sensory signals fall below the threshold of awareness (Pápai & Soto-Faraco, 2017). Thus, whether and if so, how sounds modulate the neural representation of task-irrelevant visual information is also an open question.

We addressed these questions in two different experiments: In the first experiment we tested a group of participants in a visual detection task while we concurrently registered their MEG activity. Unlike previous studies in which the effect of sounds on visual processing was characterized using univariate contrasts (Feng et al., 2014; Giard & Peronnet, 1999; Meredith & Stein, 1983; Molholm et al., 2002; Talsma et al., 2007b), here we applied an information decoding based approach. To characterize at which perceptual stages sounds modulate visual detection, we capitalized on multivariate pattern analyses to decode time-resolved d’ and c parameters from MEG activity patterns measured at different brain regions of interest (ROIs) along the visual hierarchy. Moreover, to understand whether sounds simply enhance the decodability of visual information or they also affect the maintenance of target information over time, we implemented temporal generalization (TG) analyses (King & Dehaene, 2014; Stokes, 2015).

In a second experiment we sought to understand whether sounds modulate the processing of visual stimuli automatically or through a top-down controlled mechanism. To do that, we tested a new group of participants with similar stimuli sequences as in the first experiment, but this time their attention was diverted away from the previously relevant visual gratings (Fig. 1C). Building on the same decoding methodology that we used for the first experiment, we investigated whether sounds changed the neural representations of unattended visual gratings.

**Figure 1.**
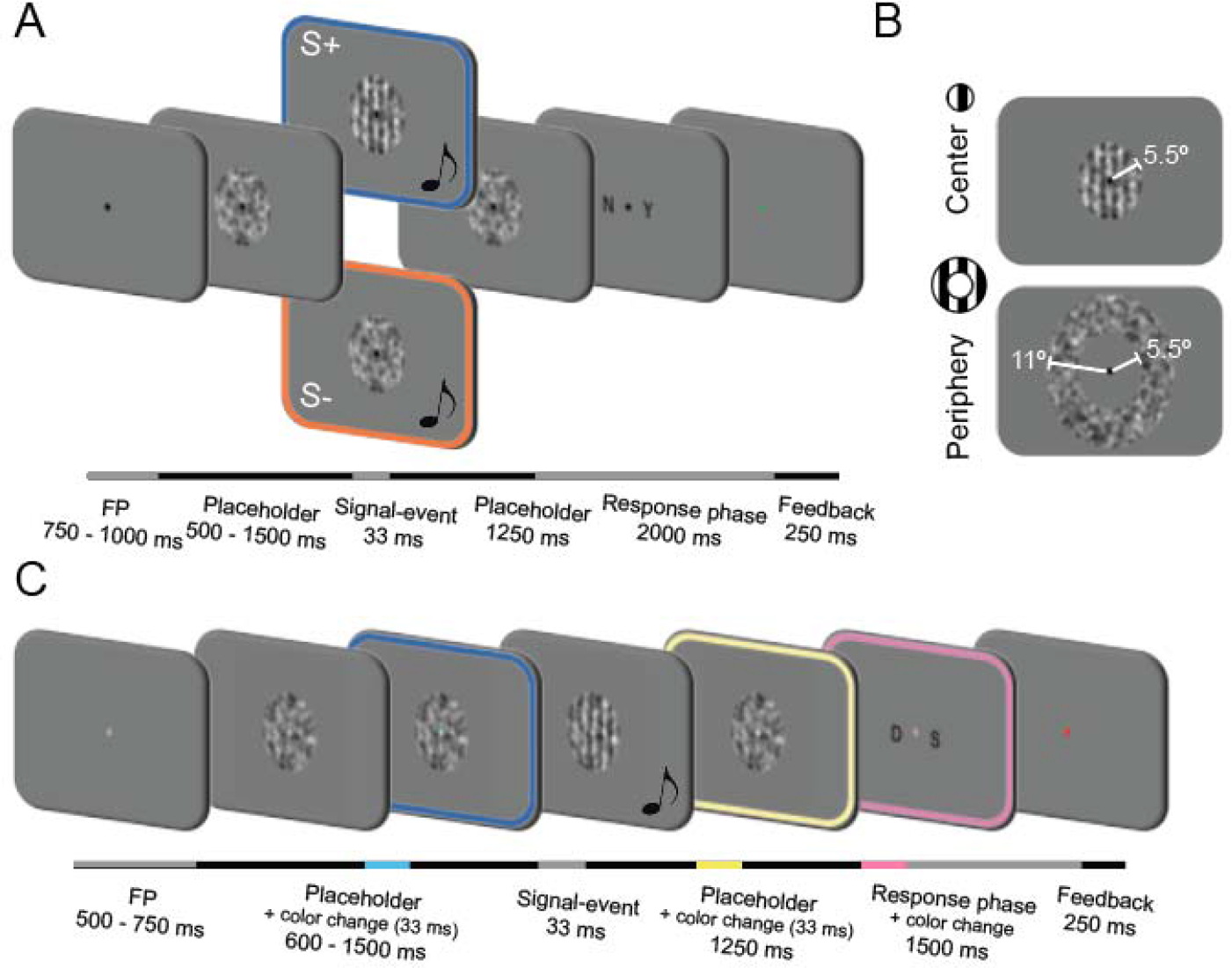
A. Experiment 1 design: Trial sequence showing an audiovisual S+ (blue) or S- (orange) event at the central visual field. In the audiovisual trials (50% of the trials), a 1000 Hz pure tone was presented simultaneously (33 ms) with the [S+ | S-] event (S+ trials = 50% and their probability of appearance was decorrelated to the sound presentation). B. Example of a central S+ and a peripheral S- stimulus. Information about their dimensions is overlaid in white. C. Experiment 2 design: The trial sequence was similar to the one used in the first experiment with the difference that now participants had to ignore the visual grating and auditory events and report on fixation point (FP) color changes. For illustrative purposes, the color frames in panel A and C highlight those events that were task-relevant in each experiment.

Foreshadowing our results below, we show that auditory input interacts with visual processing in two distinct ways: 1) Sounds enhance the decoding of the visual targets across the perceptual hierarchy, primarily in the form of an improved maintenance over time of the information encoded at post-sensory perceptual stages. This enhancement seems to be mediated by a top-down controlled mechanism, as it vanishes when the audiovisual stimuli are irrelevant for the task; and 2) Sounds automatically trigger patterns of activity in the visual cortex highly similar to the ones evoked by a visual stimulus. This result suggests that the typically reported sound-induced behavioral increment in false alarms in visual detection tasks might actually reflect a bottom-up auditory-driven perceptual bias. This is contrast to the typical interpretation of this effect as a purely decisional process (Frassinetti et al., 2002; Lippert et al., 2007; Odgaard et al., 2003).

## Results

### Experiment 1

Twenty-four participants underwent an MEG scan while they performed a visual detection task (Fig. 1A). In each trial, participants had to report the presence (S+) or absence (S-) of a briefly flashed vertical grating (33 ms). In half of the trials and orthogonal to the probability of appearance of the visual grating, an auditory stimulus was presented (the ‘audiovisual’ condition). Visual stimuli could be presented at the center (i.e. the fovea and parafovea) or the periphery (i.e. perifovea) of the visual field (Fig. 1B). This manipulation was motivated by previous neuroanatomical tracing studies in monkeys showing that more peripheral visual eccentricities receive denser projections from primary auditory cortex (Falchier et al., 2002; Rockland & Ojima, 2003), and it allowed us to explore whether the audiovisual integration strength was dependent on visual eccentricity (Fiebelkorn, Foxe, Butler, & Molholm, 2011; Gleiss & Kayser, 2013).

### Sounds improve visual detection sensitivity and reduce criterion

Observers’ sensitivity (d’) in detecting the visual target was higher in the audiovisual compared to the visual conditions (d’ = 1.57 vs 1.44: F_1,23_ = 5.67, P = 0.025, η = 0.032; Fig. 2A). This sensitivity enhancement did not depend Observers’ sensitivity (d’) in detecting the visual target was higher in the = 5.67, audiovisual compared to the visual conditions (d’ = 1.57 vs 1.44: F_1,23_L L upon visual eccentricity (interaction between modality and eccentricity: F1,23 = 0.16, P = 0.69). Furthermore, as expected, the criterion (c) parameter was lower in the audiovisual compared to the visual conditions (c = 0.56 vs 0.79: F1,23 = 23.97, P < 0.001, η = 0.014; Fig. 2B), indicating that participants had a more liberal response threshold in the audiovisual trials. Criterion did not differ between visual eccentricities (F1,23 = 0.15, P = 0.7) and the auditory-driven criterion reduction did not depend on the visual eccentricity either (F1,23 = 0.01, P = 0.9). Finally, although participants were instructed to prioritize accuracy over speed, reaction times (RTs) were faster in the audiovisual than in the visual conditions (RTs = 496 vs 540 ms: F1,23 = 61.02, P < 0.001, η = 0.06; Fig. 2C). Also for reaction times, the effect of modality did not depend upon visual eccentricity (F1,23 = 0.24, P = 0.62). Thus, our behavioral results are consistent with previous reports in showing that sounds enhance visual sensitivity, speed up visual detection and increase the proportion of reported false alarms (Fig. S2). Given that we were not able to detect any significant effect of visual eccentricity on the sound-induced visual enhancement, in the following multivariate decoding analyses the “center” and “periphery” visual field conditions will be combined to train and test the classifier. Combining these two conditions will improve the sensitivity of the classifier (as the number of trials in each training fold will be doubled). As a sanity check, we performed a control analysis where we trained and tested the classifier with data from the “center” and “periphery” conditions separately and obtained qualitatively similar results in both visual fields (Fig. S7).

**Figure 2.**
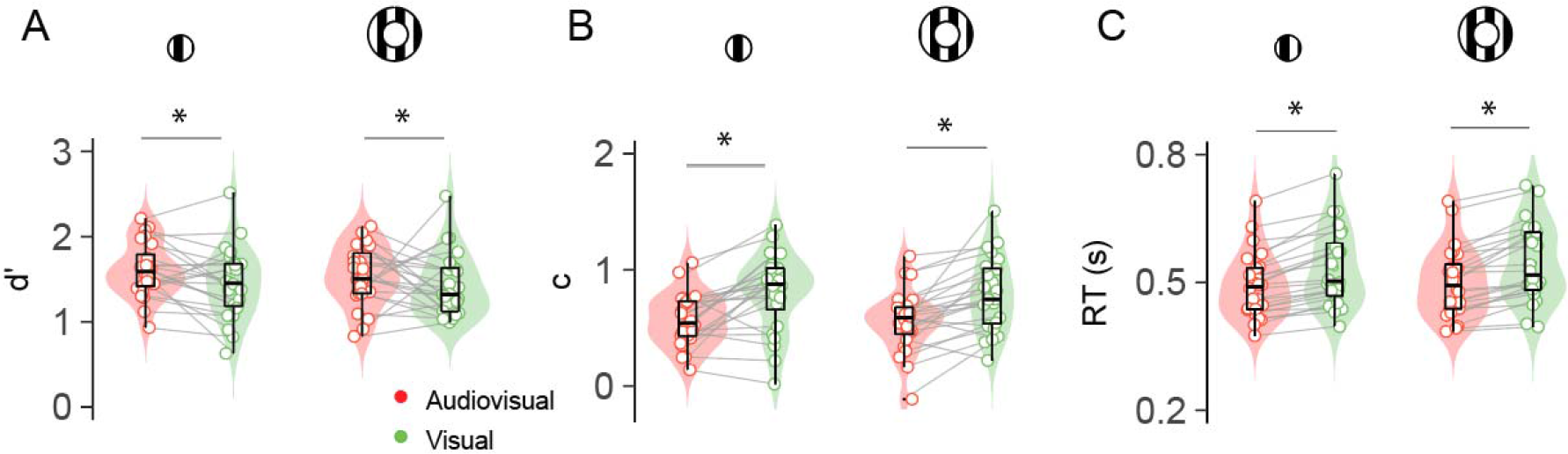
Behavioral results in the first experiment: Sensitivity (A), criterion (B) and reaction times (C) are depicted for each modality (“audiovisual”, “visual”), visual eccentricity condition (“center” and “periphery”) and participant. Inside the violin plot, the horizontal black line reflects the median, the thick box indicates quartiles, and whiskers 2 ’ the interquartile range. Grey horizontal lines connect participants scores across conditions.

### Neurally decoded signal detection parameters correlate with behaviour

To describe how sounds modulate visual processing across the perceptual hierarchy we trained a linear discriminant classifier to distinguish signal-present (S+) from signal-absent (S-) trials on the basis of MEG signals measured in visual, parietal, inferotemporal and prefrontal ROIs (upper part, Fig. 3A). These ROIs were anatomically defined and encompassed multiple sensory and decision-related brain regions involved in visual detection (King et al., 2016; van Vugt et al., 2018). Prior to testing the effect of sounds on visual detection, we explored how the neurally estimated d’ and c parameters evolved in time independent of modality (i.e., combining visual and audiovisual trials; Fig. S5 and S6).

**Figure 3.**
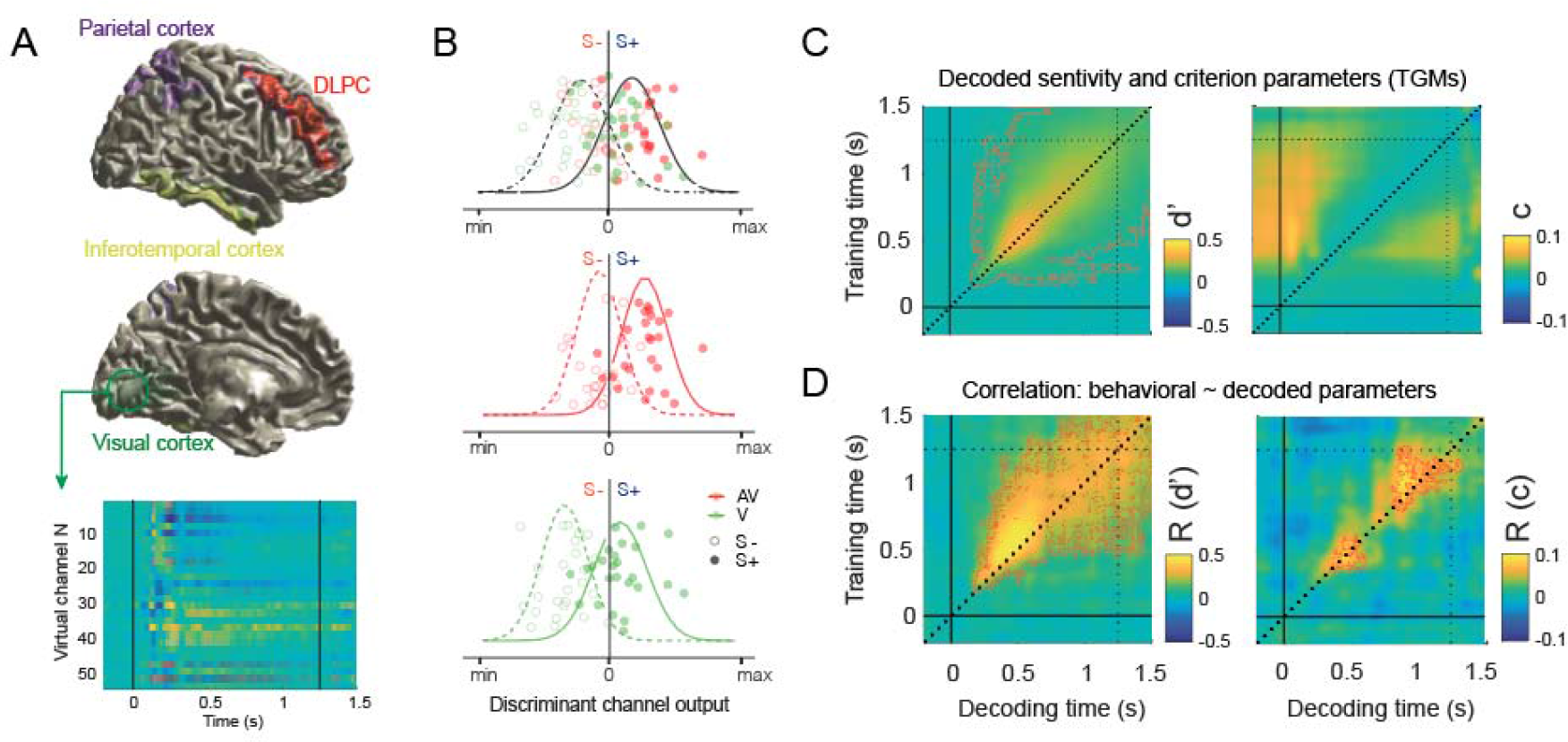
A Visual representation of the four anatomically defined regions of interest overlayed on a right hemisphere surface model of the MNI template. ROIs correspond to primary visual cortex (green), parietal cortex (purple), inferotemporal cortex (yellow) and dorsolateral prefrontal cortex (red). Source reconstructed activity extracted from the virtual channels corresponding to each ROI (e.g. visual cortex) was subsequently used to test and train a multivariate classifier (bottom part). B. A multivariate pattern classifier was trained on visual and audiovisual (green and red simulated points respectively) conditions together to optimally discriminate between S+ and S-trials (empty and full circles respectively). The trained classifier was cross validated on visual and audiovisual trials separately. We categorized S+ trials classified as signal-present as hits, and the S- trials classified as signal-present as false alarms. Note that although a classifier might be equally good in discriminating S+ from S- trials in all the conditions, there might be systematic biases to classify more often the trials in the audiovisual condition (middle panel) as signal-present than in the visual condition (bottom panel), following typical human performance. The simulated probability density functions represent how S+ (continuous) and S- (dashed) trials are distributed according to the discriminant channel output. C. Averaged over ROIs decoded sensitivity and criterion parameters (visual and audiovisual conditions together). D. For each of the six experimental blocks and for each time point combination in the temporal generalization matrix (TGM), we calculated d’ and c parameters based on the classifier output (here we show the average over the six blocks). Similarly, for each block we calculated d’ and c based on the observer’s performance. Finally, we calculated a time point by time point correlation (R) between the six neurally and behaviorally estimated d’ and c parameters. Red thin contours depict significant clusters.

We found that a classifier trained and tested on the visual and audiovisual conditions together was able to discriminate above chance level between S+ and S- trials from ∼180 ms to the end of the trial in the visual ROI (P < 0.001; see the results averaged over ROIs in Fig. 3C). A closer inspection of the d’ temporal generalization matrixes (TGMs) showed that signal decoding peaked at ∼500 ms in all the ROIs (Fig. S1.B and S5), and consistent with previous research (King et al., 2016; Mostert et al., 2015), the information represented at early sensory stages (<200 ms) generalized less strongly than at later decision-related stages. We repeated the same analysis on the neural decoder’s criterion (c). We observed that criterion increased at ∼200 ms after stimulus onset and on both sides of the diagonal (Fig. 3C). The same pattern was replicated across ROIs (Fig. S6). This result indicates that whereas a classifier trained to classify S+ and S- trials before ∼200 ms cannot be biased (given that there is not reliable information yet to be trained on), once the classifier “learns” the difference between S+ and S- trials (after ∼200 ms until the end of the trial), it will classify more often the activity in those time points not containing decodable information as S-, reflected in larger criterion values (see the results averaged over ROIs in Fig. 3D).

Next, we tested whether the neurally derived SDT parameters encoded time specific information about the participant’s latent perceptual decision-making states. We found that the neural decoder’s d’ parameters were positively correlated with the observer’s d’ along the TGM diagonal in all the ROIs, peaking at 500 ms (see ROI averaged results in Fig. 3D) and with subtle differences in the generalization spread between ROIs (Fig. S5). Further correlation analyses revealed that the neural decoder’s criterion parameters were positively correlated with the observer’s criterion parameters in an “early” cluster spanning from ∼400 ms to ∼600 ms in visual and parietal ROIs (P < 0.01), and in a “late” cluster spanning from ∼900 ms to ∼1100 ms in visual, parietal and inferotemporal ROIs (p < 0.01). These results demonstrate that the neural decoder’s d’ and c parameters are informative about participants behavior and allow us to infer at which temporal latencies the brain performs critical sensitivity and bias computations.

### Decoder’s sensitivity analysis: Sounds enhance the maintenance of postsensory visual information over time

To understand how sounds neurally enhance visual detection we contrasted the audiovisual and visual d’ TGMs in each ROI (Fig. 4A). Four significant clusters conforming a continuous pattern (P < 0.03) emerged in the inferotemporal ROI (Fig. 4B), meaning that information decodable at 500 ms was better preserved until the end of the trial in the audiovisual condition. Interestingly, in the visual cortex ROI we found that decoders trained before and after the response time (∼1100 ms to the end of the trial) and tested in the audiovisual condition were more sensitive to information present at ∼500 ms (P < 0.02) than the same decoders tested in the visual condition. This result suggests that there was an enhanced reactivation of the information encoded at ∼500 ms in the audiovisual trials, right before participants were instructed to select their final response (1250 ms). In the parietal cortex ROI we also observed a significant cluster (P < 0.01) reflecting that sounds punctually boost signal decoding at around ∼500 ms in comparison to the visual trials. This effect was time specific as it did not generalize to earlier or later time points. We registered a late but brief (∼1000 to 1100 ms) enhanced reinstatement of the information presented at ∼500 ms in the dorsolateral prefrontal cortex ROI. This small cluster (P < 0.02) preceded in time the cluster previously reported in the visual ROI (Fig. S12).

**Figure 4.**
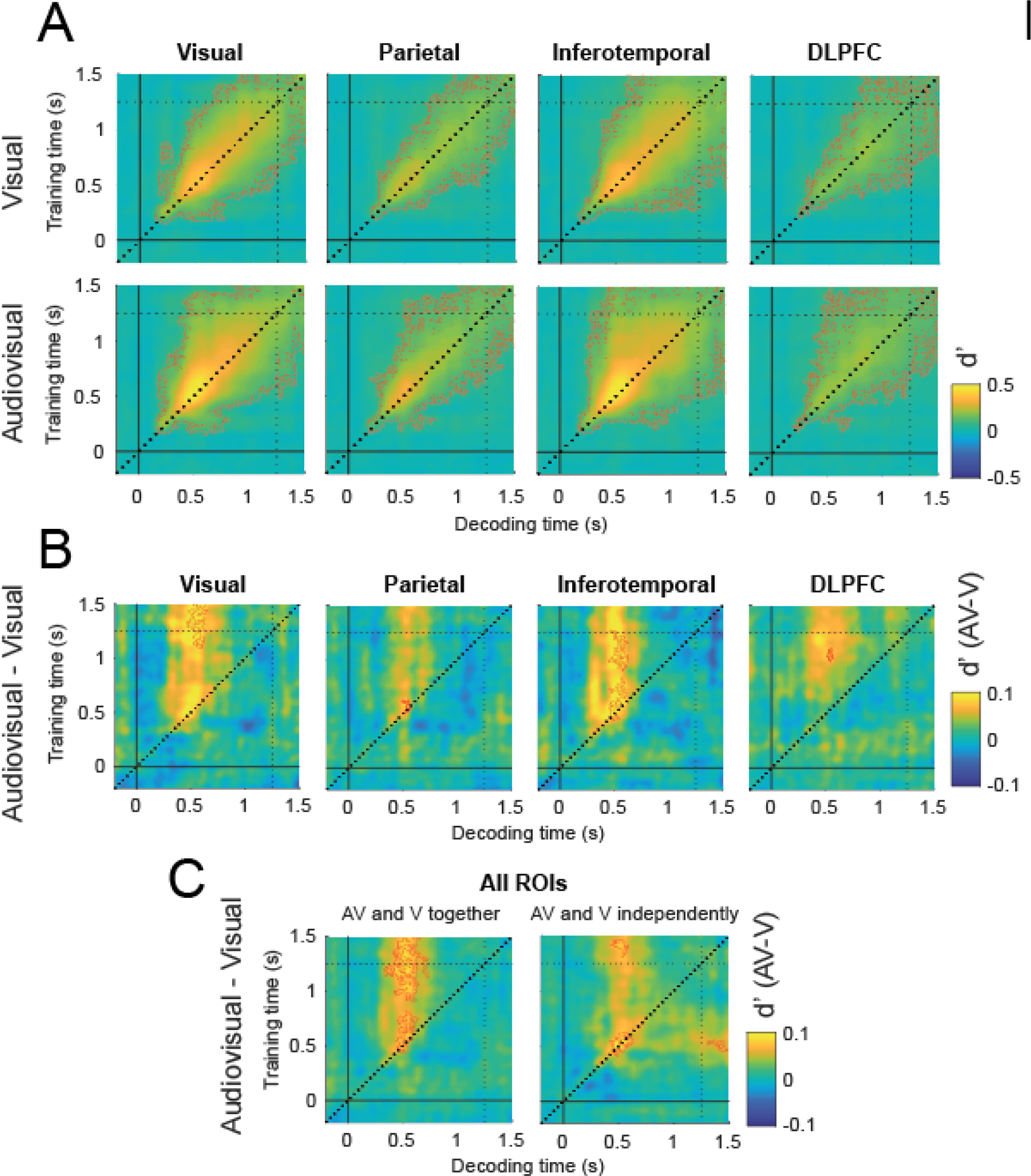
Temporal generalization matrixes depict A the neural decoder’s sensitivity parameter (d’) in the visual (upper TGMs) and audiovisual (bottom TGMs) conditions and their difference (B) in each ROI. C represents the differential decoded sensitivity averaged over ROIs with classifiers trained in the visual and audiovisual conditions together (left panel) or separately (right panel). Independently trained classifiers replicate similar results as when the classifiers were trained in both conditions together and recovers the expected decoding symmetry with respect to the diagonal. Red contours delimit significant clusters.

To obtain a general understanding of the TG pattern, we looked at the d’ TGM averaged over ROIs. Consistently with the individual ROIs decoding analyses we found that the audiovisual enhancement of information conformed a continuous pattern (Fig. 4C). Decoders trained from ∼500 ms to the end of the trial could decode better the visual stimulus at 500 ms in the audiovisual compared to the visual condition. This vertical pattern was composed by two elongated clusters (P < 0.001 and P < 0.005) separated by a small gap spanning from approximately 750 to 950 ms where the sound-induced visual decoding enhancement was not significant.

Surprisingly, we found that although multiple decoders trained at different time points could decode the signal presence at 500 ms, a decoder trained at 500 ms was not able to “symmetrically” enhance the decoding of the audiovisual trials within an analogous temporal window. In a complementary control analysis (Fig. 4C and Fig. S8), we confirmed that by training and testing the classifier in the visual and audiovisual conditions separately, the sound-induced d’ enhancement pattern became symmetrical with respect to the diagonal. This result indicates that training the classifier in the visual and audiovisual conditions together might come with the cost of losing/averaging out the idiosyncratic features that characterize the visual and audiovisual neural activity patterns. Therefore, a classifier trained on data from both conditions together at the decoding peak (500 ms), but tested in the visual and audiovisual conditions separately might have reduced capacity to generalize the information and tease apart between S+ and S- trials when the most discriminative information decays (i.e. before and after the decoding peak), and other more modality-specific neural modulations take over.

In summary, our d’ analyses show that sounds enhance the maintenance over time of late perceptual information (500 ms) until the response phase, across multiple ROIs.

### Decoder’s criterion analysis: Neurally decoded criterion is lower in the audiovisual trials

Consistent with participants performance, we found that the neural decoder’s criterion was reduced in the audiovisual compared to the visual condition in all the ROIs (Fig. 5A). That is, a decoder trained on visual and audiovisual trials together systematically classified more often the audiovisual trials as S+ trials than the visual ones (Fig. S4). This pattern was predominant along the TGM diagonals with some qualitative differences between ROIs (Fig. 5B). In visual and inferotemporal cortex ROIs we observed significantly negative clusters (P < 0.005), indicating that multiple decoders trained from ∼500 to 1100 ms had a generalized bias to classify audiovisual trials patterns occurring at ∼400 ms as S+ trials. In the parietal and dorsolateral prefrontal ROIs, a similar but less generalized pattern emerged along the diagonal (P < 0.01). In addition, we observed a second significant cluster that was specific to the parietal and dorsolateral ROIs. In this second cluster, the direction of the bias reversed. That is, a decoder trained around ∼400 ms was biased to classify more often a trial as S+ in the visual than in the audiovisual trials (from ∼600 to 1100 ms in the parietal ROI; P < 0.03, and from ∼1000 to 1100 ms in the dorsolateral ROI; P < 0.02). The latency of this positive bias may indicate that whereas in the S- audiovisual trials the participants “believed” more consistently to have seen the target immediately after the sound presentation, in the visual trials, due to the absence of an auditory cue signalling the most likely latency of the visual stimulus onset, the belief of seeing the target can potentially spread to later time points.

**Figure 5.**
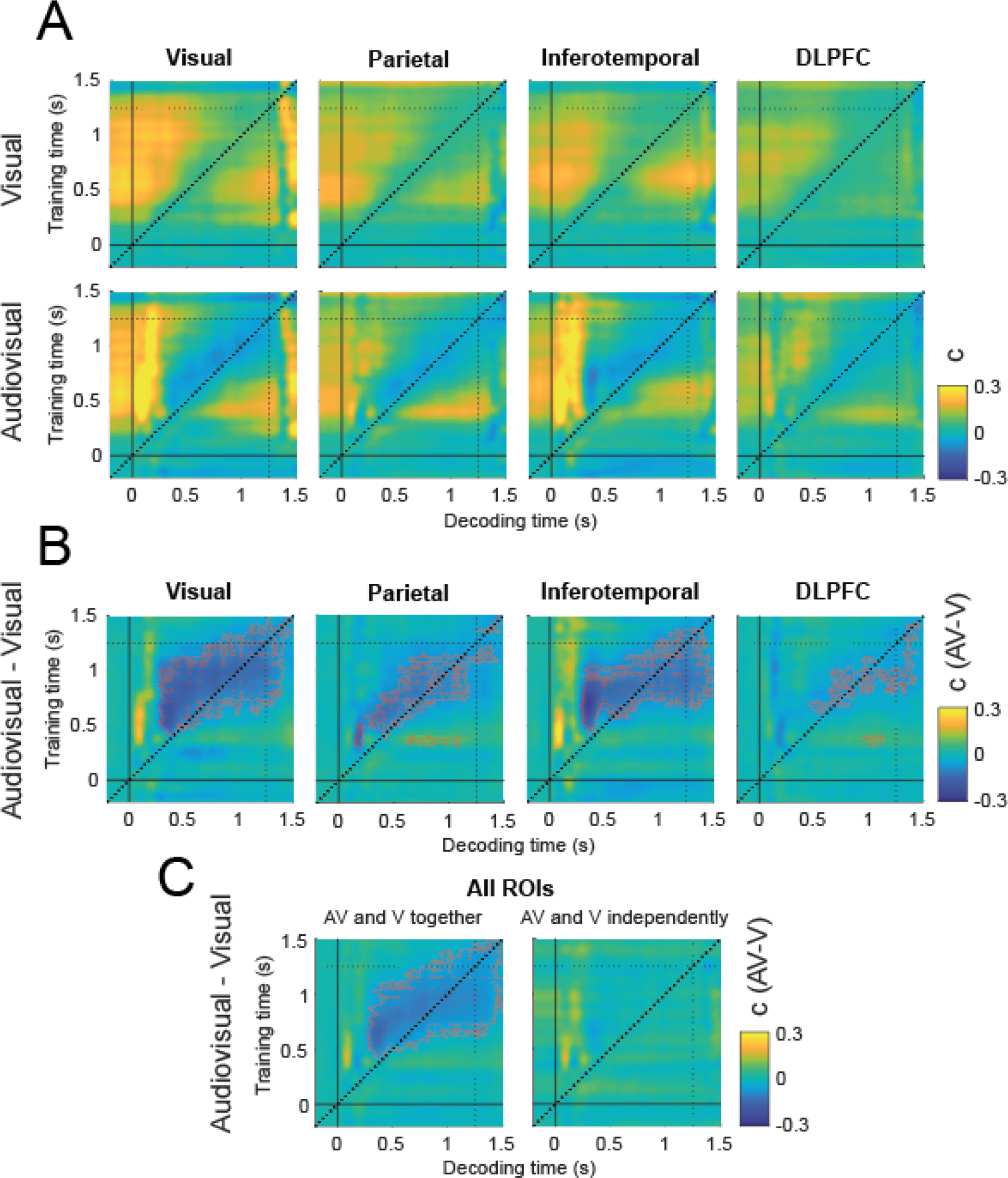
Temporal generalization matrixes depict A. the neural decoder’s criterion parameter (c) in the visual (upper TGMs) and audiovisual (bottom TGMs) conditions and their difference (B) in each ROI. C represents the differential decoded criterion averaged over ROIs with classifiers trained in the visual and audiovisual conditions together (left panel) or separately (right panel). Independently trained classifiers fail to capture the existing differences in criterion between the audiovisual and visual conditions. Red contours delimit significant clusters.

In summary, criterion analyses demonstrate that sounds evoke neural activations that are more often misclassified as S+ trials than in the visual conditions. As expected, in a control analysis we showed that this sound-induced criterion bias would be inappreciable if we would have trained and tested the decoders in the visual and audiovisual conditions separately (Fig. 5C).

### Experiment 2

In our second experiment we tested if the mechanism by which sounds enhanced the maintenance of visual information is automatic and bottom-up driven or requires participants to be actively engaged in a visual detection task. For instance, it is possible that subjects explicitly used the auditory stimulus as a cue to optimally orient their attention in time, prioritizing the encoding and maintenance of the most informative visual samples in short-term memory (Lippert et al., 2007; Los & van der Burg, 2013; Sergent et al., 2013). Such a mechanism cannot be considered bottom-up as it requires an endogenous control of attention. Relatedly, we also investigated whether the reduced criterion in the audiovisual condition reflects a decisional or a perceptual bias.

To address these questions, a new group of participants (n = 24) was presented with similar stimuli sequences as in our first experiment. However, participants now had to attend to and memorize a sequence of three fixation point color changes (Fig. 1C), rendering the previously relevant vertical gratings and auditory stimuli task irrelevant. The unattended gratings were presented at only one visual eccentricity (“central”) and with two different levels of contrast (i.e. contrast was set at threshold level in the low contrast “S+” condition and at two-times the threshold level in the high contrast “S++” condition). Adding this new high contrast condition allowed us to test how perceptually salient but ignored S++ stimuli interact with sounds, and ensured that unattended visual stimuli evoked sufficient signal to be decoded from neural activity patterns.

We hypothesised that 1) if the enhanced maintenance of neurally decoded visual information described in our first experiment is supported by automatic bottom-up multisensory integration, we should replicate the same result in this second experiment despite participants ignore the audiovisual stimuli. Following up on the same logic, 2) if the reduction in the decoder’s criterion following the sound presentation described in the first experiment corresponds to a decision-level bias, we should not be able to replicate here the same result as participants do not make decisions about the visual gratings. Instead, finding again a more biased decoder in the audiovisual compared to the visual condition would imply that sounds can automatically trigger patterns of activity in the brain that are misclassified by the decoder as a visual stimulus.

### Audiovisual distractors do not interfere with participants performance in the working memory task

Participants’ performance in remembering the sequence of fixation point colour changes was high (group mean d’ = 2.99, SD = 0.91). Importantly, their sensitivity and criterion parameters did not vary as a function of gratings levels of contrast (d’ F2,48 = 0.98, P = 0.37 and c F2,48 = 0.29, P = 0.75) or auditory stimulus presentation (d’ F1,24 = 0.308, P = 0.58 and c F1,24 = 0.44, P = 0.51; Fig. S2). This demonstrates that the task-irrelevant audiovisual stimuli could be successfully ignored by the participants as they did not interfere with the color working-memory task. Likewise, RTs analyses did not show any difference as a function of grating contrast (F2,48 = 0.9, P = 0.41) or auditory presentation (F1,24 = 0.01, P = 0.89. Group mean RT = 880 ms, SD = 0.15).

### Decoder’s sensitivity analysis: Sounds do not automatically enhance the maintenance of visual information

We found that the decoders trained and tested in the low contrast condition could not discriminate between the S+ and S- trials (Fig. S9 and S10). This result indicates that diverting participants attention away from the visual gratings diminished the contrast response gain (Herrmann et al., 2012) compared to the first experiment, leading to a null decoding sensitivity for threshold-level visual stimuli. On the other hand, the neural activity driven by the high contrast stimuli was strong enough to be discriminated above chance level at multiple time points (Fig. 7). Given the null sensitivity of the decoders in classifying the neural activity patterns associated to low contrast stimuli, we constrained our subsequent analyses to the high contrast condition.

We found that high contrast stimuli could be decoded as early as ∼100 ms in the visual cortex. Decoding accuracy peaked at 180 ms in the visual, inferotemporal and parietal ROIs and around 350 ms in the dorsolateral prefrontal ROI. The d’ TGM profile in the high contrast condition showed that most of the significant clusters were distributed along the diagonal with weak information generalization (Fig. 6A). This is consistent with the highly dynamical information broadcasting that takes place during sensory processing (King et al., 2016; Mostert et al., 2015), and suggest that the stimuli were ignored by the participants as they failed to trigger the sustained generalization pattern typically associated with visual awareness (King et al., 2016).

**Figure 6.**
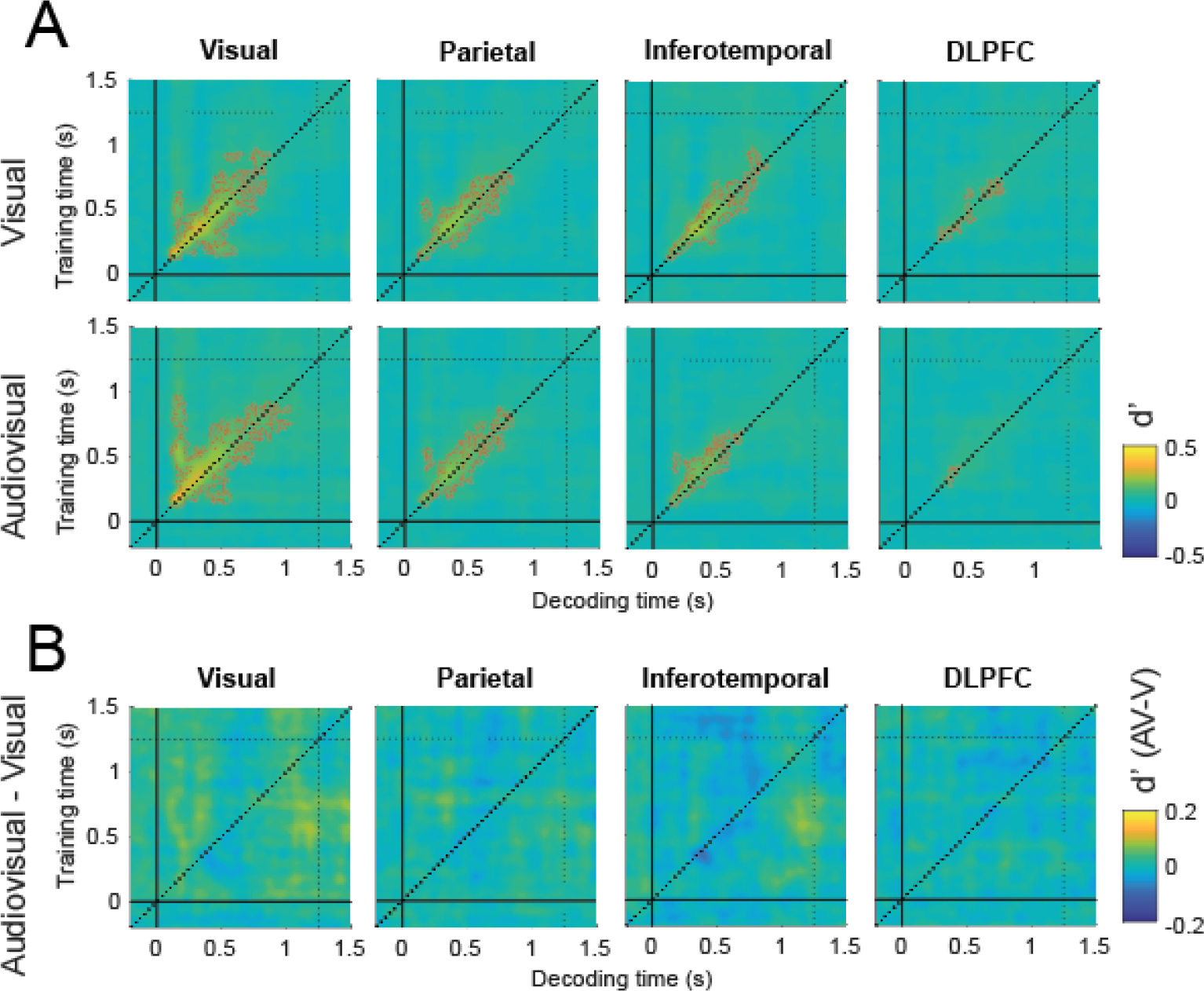
Temporal generalization matrixes illustrating decoded sensitivity parameters (d’) in the high contrast condition (S- vs S++). A and B follow the same labelling convention as in figure 5.

To investigate whether sounds enhanced the neural decodability of the unattended visual targets, we contrasted the audiovisual and visual d’ TGMs but we could not find significant differences between both modality conditions in any of the selected ROIs (Fig. 6B). Since this contrast yielded clear differences in our first experiment, but we observe a null result here when attention is directed away from the stimuli, we conclude that the sound-induced enhanced maintenance of visual information, reflected in sensitivity, is highly dependent on stimulus relevance, and it is therefore likely mediated via top-down mechanisms.

### Decoder’s criterion analysis: Sounds drive visual activity patterns in visual cortex in a bottom-up fashion

To test whether sounds influenced the neural decoder’s criterion in the high contrast condition (Fig. 7A), we contrasted the audiovisual and visual c TGMs. We found a significant cluster (P < 0.01) in the visual cortex ROI, meaning that those decoders trained from ∼450 to 650 ms classified more often the activity patterns from ∼250 to 450 ms as S+ in the audiovisual compared to the visual trials (Fig. 7B). That is, sounds increased the proportion of neurally estimated hits but also of false alarms at around 350 ms (Fig. S5). This result demonstrates that sounds can automatically evoke patterns of activity in the visual cortex that are often misclassified by a decoder as an actual visual stimulus.

**Figure 7.**
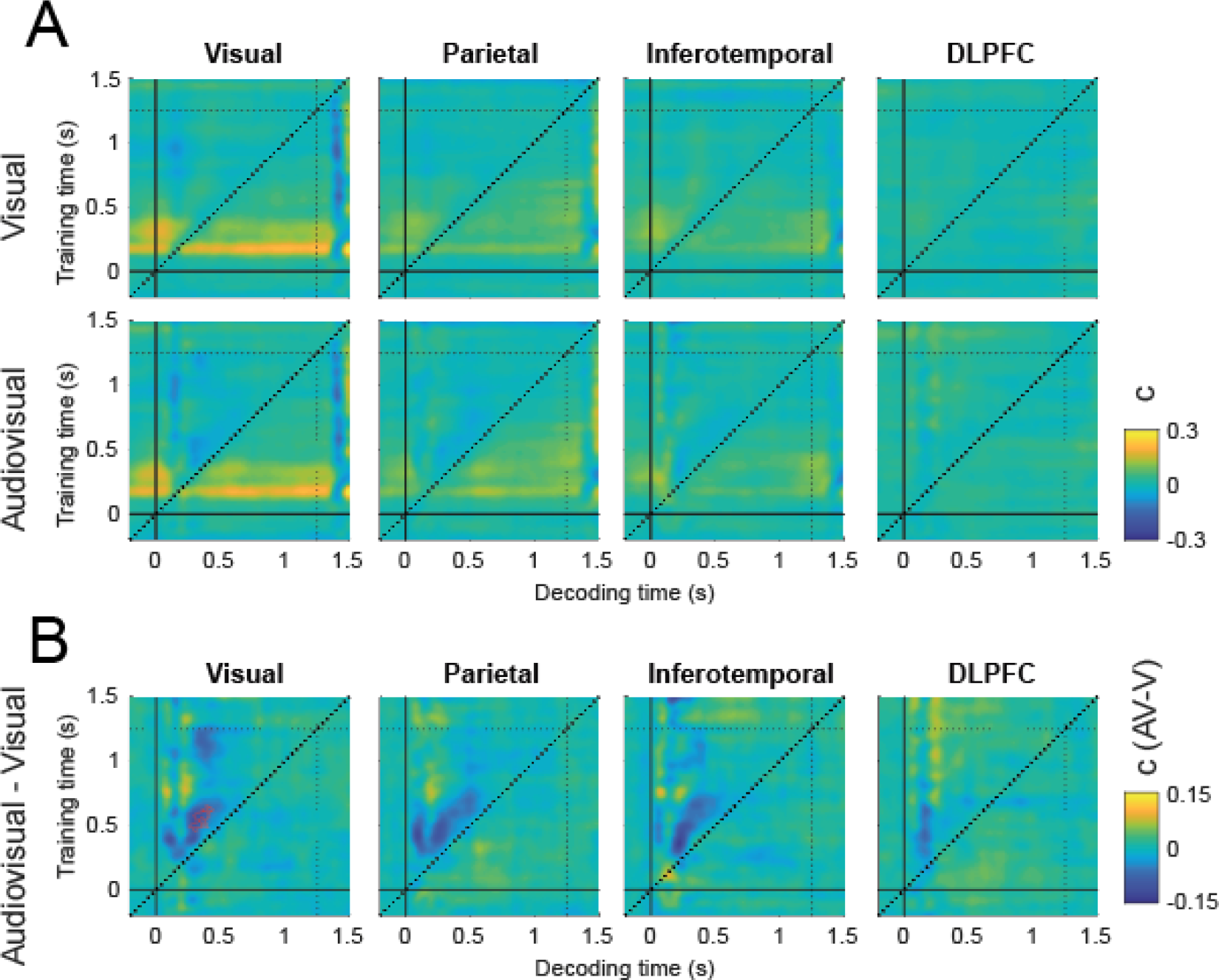
Temporal generalization matrixes generated from decoded criterion parameters (c) in the high contrast condition (S- vs S++). A and B follow the same labelling convention as in figure 5.

One might argue that the sound-induced increase in false alarms reports might be unspecifically related to the decoder picking up on univariate auditory event-related field (ERF) deflections present in both S+ and S- trials. This explanation is unlikely because the decoders that showed an increase in false alarms were trained at a temporal window (450 to 650 ms) in which the neural activity in S- and S+ trials did not differ univariately (Fig. S11). Moreover, the activity that was misclassified as S+ spanned along the second half of the auditory ERF but did not generalize to the first half even though both halves were almost symmetrical. Our results are congruent with previous literature in showing that sounds activate early visual regions in a bottom-up fashion (Deneux et al., 2019; Feng et al., 2014; Ibrahim et al., 2016; Romei et al., 2009b), and suggest that these activations encode stimulus-specific neural representations.

## Discussion

In our first experiment we used a multivariate decoding approach to dissociate the neural mechanisms by which sounds modulate observers’ sensitivity and bias in a visual detection task. Under the assumption that sounds can affect the decoding amplitude but also the latency or duration of visual representations, we applied TG analyses (King & Dehaene, 2014; Stokes, 2015) revealing that indeed sounds enhanced the maintenance of visual information over time. The effect of sounds on visual sensitivity can be characterized by different dynamical processing models (see models’ description in Fig. 8) that vary in terms of their overall architecture (i.e., the number and the order of the processing stages) and whether they postulate that the sound-induced enhancement correlates with an increase of the amplitude or the duration of a given processing stage (Gwilliams & King, 2020; King et al., 2016).

**Figure 8.**
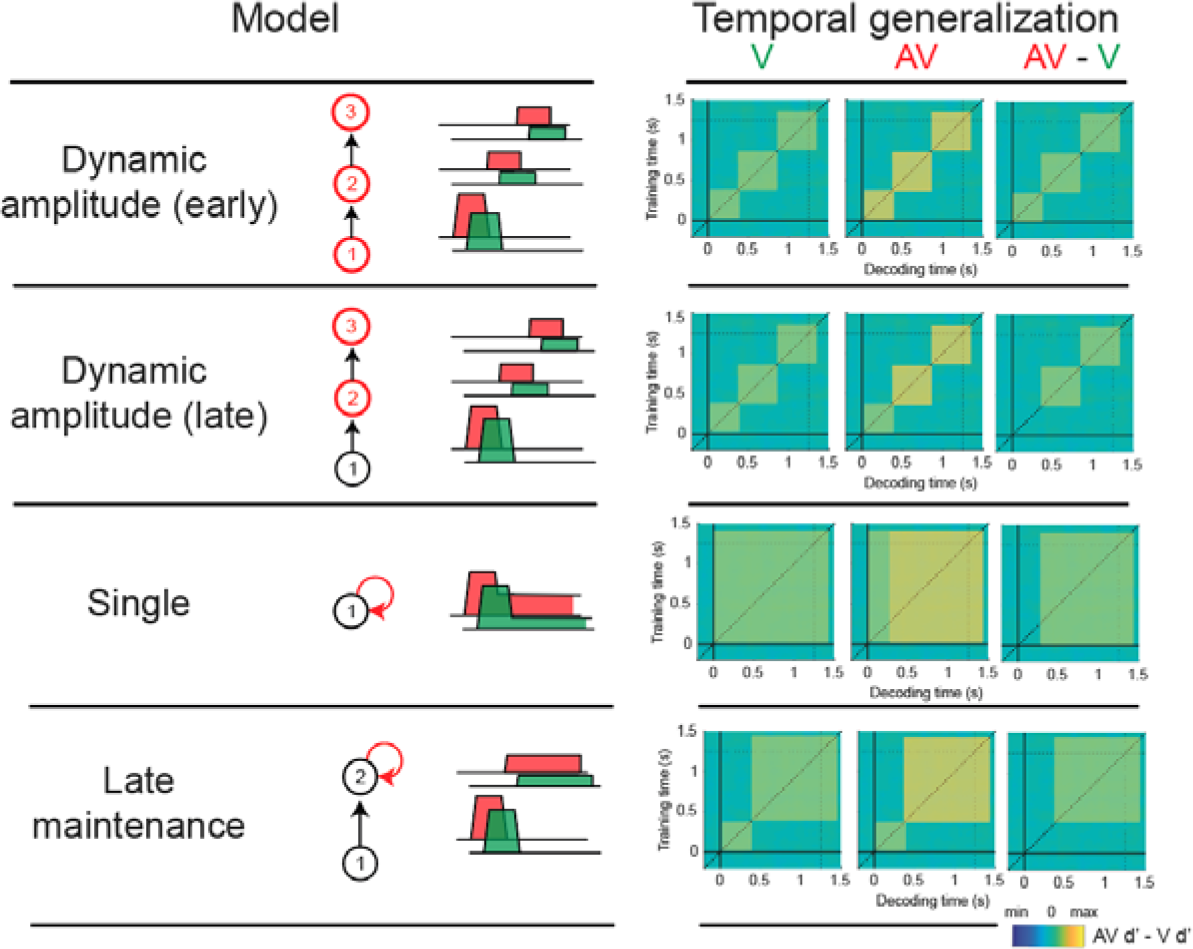
Sounds might improve the detectability of a visual signal through different mechanisms. To characterize the dynamic nature of information processing we describe four models that vary in terms of the number and ordering of processing stages. We propose that sounds could enhance detection by increasing 1) the decoding amplitude or 2) the duration of one or several processing stages: The “dynamic amplitude” models broadcast target information across a sequence of short-lived processing stages whose amplitude codes for its detectability. We hypothesize that sounds might act upon one of these stages, boosting the decoding amplitude at early or late processing stages. The “single stage” model encodes and maintains the target information within the same processing stage. We propose that sounds might improve the maintenance of information within the initial processing stage. The “late maintenance” model transmits the encoded information to a later processing stage where task-relevant information is sustained through recurrent processing loops. Sounds might enhance the maintenance of information within the later processing stage. Vertical arrows represent information transfer from one neural stage to the next. Circular arrows represent recurrent feedback loops that maintain information over time. The key process that characterizes the effect of sounds in each model is highlighted in red.

Our results are better explained by a “late maintenance” model in which stimulus information after being encoded in early sensory regions is remapped as a decision variable (DV) and stored until the response phase. According to this model’s predictions, sounds would improve visual sensitivity by enhancing the maintenance over time of task-relevant visual information encoded at ∼500 ms after the stimulus onset (see that the target information encoded at 500 ms was highly correlated with participants sensitivity). Based on the temporal ordering of the significant clusters across ROIs (Fig. S12) and in line with the currently accepted view that multisensory interactions depend upon a widely distributed network of brain regions (Cao et al., 2019; Rohe et al., 2019; Rohe & Noppeney, 2015, 2016), we propose that a late maintenance dynamical mechanism could be articulated, first by the encoding of the visual stimulus in the visual cortex. Concurrently, the encoded sensory information would be mapped into a latent DV in parietal regions (Kiani & Shadlen, 2009; O’Connell et al., 2012; Zhou & Freedman, 2019). This is a stochastic process in which an auditory cue might amplify the participant’s gain towards those DV samples that more likely encode the visual target. These highly informative samples could be stored in the inferotemporal and dorsolateral prefrontal cortex to protect them from interference with new incoming sensory input. Indeed, the interplay between these two brain regions has been demonstrated to play an important role in short-term memory during perceptual decisions (D’Esposito & Postle, 2015; Gross et al., 1972; Pagan et al., 2013; Stokes et al., 2013; van Vugt et al., 2018). As the response phase approaches, the participants can strategically reorient their featured-based attention towards the stored relevant stimulus information. This manifests by an enhanced reactivation of the target in the DLPF cortex (Squire et al., 2013), that subsequently leads to an enhanced reinstatement of the stimulus information in early visual regions (Christophel et al., 2017; D’Esposito & Postle, 2015; Sprague et al., 2016).

We could not replicate the same pattern of results in the second experiment when participants ignored the audiovisual stimuli. This suggests that in the first experiment sounds helped participants to guide their temporal attention in a top-down fashion, improving the processing and maintenance of task-relevant perceptual information (Lippert et al., 2007; Los & van der Burg, 2013; Ngo & Spence, 2012). We speculate that the sound-induced visual sensitivity enhancement could be partly mediated by a mechanism like retroception (Sergent et al., 2011, 2013). In retroception a post-stimulus cue can retrospectively amplify the signal gain of a target stored in perceptual memory that would otherwise have escaped consciousness. In the sound-induced visual enhancement such signal gain amplification would be instantiated by an auditory cue instead, improving the maintenance of perceptual information stored at post-sensory (i.e. decision) processing stages.

We must acknowledge that showing that the crossmodal visual enhancement takes place through a top-down controlled mechanism does not preclude that other more automatic and sensory-level multisensory mechanisms could contribute in parallel to enhance visual sensitivity (although these might be out of reach of our decoding analyses sensitivity). For instance, audiovisual inputs might be integrated at subcortical level in the superior colliculus or through direct connections between early sensory areas (Clavagnier et al., 2004; Deneux et al., 2019; Garner & Keller, 2021; Ibrahim et al., 2016; C. Kayser & Logothetis, 2007; Meredith & Stein, 1983). Alternative accounts propose that sounds could exert a modulatory influence by resetting the phase of ongoing oscillations in visual cortex, such that co-occurring visual targets align with high-excitability “ideal” phases (Fiebelkorn, Foxe, Butler, Mercier, et al., 2011; Lakatos et al., 2007; van Atteveldt et al., 2014). These two sensory-level mechanisms of audiovisual integration have been well-described at an electrophysiological level, however future studies must continue elucidating which are their specific functional role in multisensory perception.

In this work, in addition to the sound-induced enhancement in visual sensitivity, we showed that sounds reduced the participants criterion in visual detection. Consistently with this result, in our first experiment the neural decoding analyses showed that sounds biased the decoders to classify more often the audiovisual trials as S+ in multiple time points. The classical interpretation of this decoding bias implies that participants, after hearing the sound, were more inclined to believe that the target has been presented (i.e. decision-level bias). However, in our second experiment in which participants did not have to make any decision about the presence or absence of the visual targets, we showed that sounds automatically evoked patterns of activity in early visual regions similar to the patterns evoked by an actual visual stimulus. This sound-induced visual activity patterns could be subjectively misperceived by an observer as a visual stimulus, leading to perceptual-level biases.

This result is consistent with previous work (Ibrahim et al., 2016; Romei et al., 2007, 2009b, 2012) by demonstrating that sounds alone can modulate the activity in early visual areas in a bottom-up fashion. It is possible that an abrupt sound that has been repeatedly paired with a visual stimulus, when presented in isolation, may automatically reactivate the associated (visual) sensory traces (den Ouden et al., 2009; Garner & Keller, 2021). Indeed, this mechanism might be at the base of many multisensory illusions like the double-flash illusion in which pairing two auditory stimuli with one visual stimulus induces the percept an additional illusory visual stimulus with identical physical properties as the inducer stimulus (Berger et al., 2003; McCormick & Mamassian, 2008; Pérez-Bellido et al., 2015). In brief, our results provide neural evidence for an alternative hypothesis on the interpretation of SDT parameters (J. Witt et al., 2012; J. K. Witt et al., 2015) that challenges the classical assumption postulating that sound-induced reductions in criterion necessarily correspond to decision-level biases.

Retrograde tracing studies in monkeys have revealed the existence of direct connections between the auditory the visual cortex. These connections are heterogeneously distributed across the visual field with more peripheral eccentricities receiving denser projections (Clavagnier et al., 2004; Rockland & Ojima, 2003). In our first experiment, we explored whether such asymmetrical connectivity pattern translated into different multisensory interactions. To do that, we included two conditions in which the visual stimuli could appear at the “center” or the “periphery” of the visual field. However, our behavioral and neural decoding analyses yield similar results in both eccentricity conditions. Future studies should further investigate whether the heterogenous auditory-to-visual cortex connectivity pattern reported in monkeys replicates in humans (Beer, Plank and Greenlee, 2011), and if it actually involves a functional dissociation for multisensory processing.

## Conclusion

In recent years it has been demonstrated that the visual and auditory systems are heavily interconnected at multiple processing levels (Driver & Noesselt, 2008; Murray et al., 2016; Rohe & Noppeney, 2015). Although the effect of sounds on visual detection has been well described at behavioural level, it remained unclear which specific neural mechanisms give support to crossmodal interactions. In this study we devised a novel approach to neurally dissociate the contribution of sounds to sensitivity and criterion modulations in visual detection tasks. Our results demonstrate that multisensory interplay in visual detection does not exclusively rely on sensory-level crossmodal interaction. Instead, it unfolds at multiple levels of the perceptual hierarchy improving the amplitude of the encoded visual representations and their temporal stability. In addition, our results also help to reconcile two opposing views by showing that audiovisual interactions in detection rely on parallel top-down and bottom-up crossmodal mechanisms. Whereas the sound-induced improvement in visual sensitivity is mediated through a widely distributed network of brain regions in which the maintenance of post-sensory visual information is improved via top-down mechanism, sound-induced reductions in criterion are primarily reflected in bottom-up sound-driven modulations of early visual cortex activity. In the future, the use of temporal generalization decoding in combination with more specific functional and anatomical localizers will help to understand with a greater level of detail which specific information is stored in each processing stage and how it is transformed across the multiple stages that conform the perceptual decision-making process.

## Materials and Methods

### Subjects

Twenty-five healthy human volunteers with normal or corrected-to-normal vision and audition participated in the first t (17 females, mean age = 24 years, SD = 6 years) and second experiment (12 females, mean age = 25 years, SD = 7 years).

One subject in the first and one subject in the second experiment were excluded during the preprocessing due to insufficient data quality (severe eye and muscle artifacts and poor performance). The sample size was determined prior to data collection, and ensured 80% power to detect medium-to-large effects (Cohen’s d>0.6). Participants received either monetary compensation or study credits. The study was approved by the local ethics committee (CMO Arnhem-Nijmegen, Radboud University Medical Center) under the general ethics approval (“Imaging Human Cognition”, CMO 2014/288), and the experiment was conducted in compliance with these guidelines. Written informed consent was obtained from each individual prior to the beginning of the experiment.

### Stimuli

Visual stimuli were back-projected onto a plexiglass screen using a PROPixx projector at 120 Hz. In the first experiment the screen region where the targets could appear was delimited at the beginning of each trial using parafoveal (“center” condition; inner and outer perimeters 1° and 5.5° radius) or perifoveal annular (“periphery” condition; inner and outer perimeters 5.5° and 11° radius respectively) noise patches centered on the fixation point (Fig 1B). In the second experiment all the stimuli were presented parafoveally (“center”). The noise patches were created by smoothing pixel-by-pixel Gaussian noise through a 2D Gaussian smoothing filter (Wyart et al., 2012). Signal-present (S+) stimuli consisted of vertical sinusoidal gratings (spatial frequency of 0.5 cycles/° and random phase sampled from a uniform distribution) added to the previously generated noise patches. Thereby, the noise structure of the placeholder and signal stimuli was the same within trials, but it was randomly generated for each new trial. The fixation point was a circle (radius = 1°) presented at the center of screen. In the first experiment the fixation point color was black and in the second experiment light grey (luminance: 405 cd/m^2^). All the visual stimuli were displayed on a gray background (50% of maximum pixel intensity, luminance: 321 cd/m^2^). Auditory stimuli were presented through in-ear air conducting MEG compatible headphones and consisted of binaurally delivered pure tones at 1000Hz (∼70 dB, 5 ms rise/fall to avoid clicks; 16 bit mono; 44.100 Hz digitization). All the stimuli were generated and presented using MATLAB (The Mathworks, Inc., Natick, Massachusetts, United States) and the Psychophysics Toolbox extensions (Brainard, 1997).

### Procedure and experimental design

In the first experiment participants performed a visual detection task (Fig. 1A). First, the participants completed a 5 minutes behavioral training session in which they were familiarized with the new environment and task. After the practice session, we used an adaptive staircase (Quest; Watson & Pelli, 1983) to estimate the level of contrast at which each subject correctly detected a vertical grating in 70% of the cases. The trial sequence used during the Quest procedure was similar to the one used during the main task detection blocks (see below) except that subjects were exclusively presented with the visual condition. We used two independent staircases to estimate the contrast for the center and periphery conditions and only the trials in which the grating was presented were used to update the staircase. Once we estimated the participant threshold, participants started the main task blocks. The appearance of a central fixation point signalled the beginning of each trial. This fixation point was kept on the screen for a variable period of time of 750 to 1000 ms (randomly drawn from a uniform distribution) and determined the inter-trial-interval (ITI). Then, a circular or an annular noise placeholder was displayed on the screen. The noise placeholder constrained the visual space prior to the presentation of the visual target indicating at which visual eccentricity the target might occur (“center” or “periphery”, both conditions were equiprobable) and eliminating any possible spatial uncertainty. After a random period of time between 1000 to 1500 ms, the target grating, referred to as signal-present trial (S+) or the same noise placeholder, referred to as signal-absent trial (S-) was displayed for 33 ms (4 frames). Importantly, in half of the trials an auditory tone of the same duration as the target was presented in synchrony with the [S+ | S-] event onset. Subsequently, the noise placeholder alone was displayed again and remained on the screen for a fixed period of 1250 ms. Then, the letters ‘Y’ and ‘N’ (as abbreviations for “Yes” and “No”, respectively) were centered around 4° the fixation dot. Subjects reported their decision as to whether or not they had detected a vertical grating by pressing a button with either the left or the right-hand thumb, corresponding to the position of the letter that matched their decision. The position of the letters (‘Y’ left and ‘N’ right, or ‘N’ left and ‘Y’ right) was randomized across trials to orthogonalize perceptual decision and motor response preparation. Finally, after a response period of 2000 ms the fixation point turned green, red or white for 250 ms signaling “correct”, “incorrect” or “non-registered” responses respectively. Participants performed a total of 576 trials divided in 6 blocks of ∼10 minutes. There were a total of 8 different experimental conditions: stimulus type (“signal-present” / “signal-absent”) × modality (“visual” / “audiovisual”) × eccentricity (“center” / “periphery”), and participants ran a total of 72 trials per combination of conditions. During the main task the grating contrast values were kept fixed within blocks. If at the end of one block, due to learning or tiredness participants showed near perfect (>90% correct responses) or chance level (<55% correct responses) performance, to prevent ceiling or floor effects the target contrast was updated before the beginning of the next block using the new threshold value estimated by the Quest.

In the second experiment we designed a task that forced participants to ignore the previously task-relevant visual gratings and sounds, and test whether the effects found in the first experiment were simply due to participants being involved in a visual detection task. Participants completed a 5 minutes behavioral training session in which they were familiarized with the task (see below). After the practice session, as in the first experiment we used a Quest to estimate the level of contrast at which each subject correctly detected a vertical grating in 70% of the cases. Then, after ensuring that participants had understood the main task, we continued with the experiment. In the main tasks blocks the stimulus sequence was identical to the one used in the first experiment with some minor but relevant modifications (Fig. 1C). Here, although the vertical gratings and sounds were presented in each trial as in the first experiment, the participant’s task was to ignore them and perform a working memory task on the fixation point: The fixation point changed to magenta, yellow or cyan briefly for 32 ms once before and once after the ignored [S+ | S-] event and participants had to memorize these two color changes. At the end of the trial and during the response phase, the fixation point changed a third time revealing a color that could be the same or different (in 50% of the trials) as one of the previously memorized colors. Participants had to choose whether this third color was repeated or not by selecting between ‘S’ or ‘D’ (as abbreviations for “Same” and “Different” respectively) using the left or right thumb on the button box. The fixation point color sequences were generated in each trial by randomly selecting without repetition two of the three previously described colors. To ensure that participants maintained their attention steadily on the fixation point during the whole trial, the first two fixation point color changes happened during the presentation of the first and second noise placeholders, and their onset varied randomly from trial to trial 300 to 750 ms relative to the onset of the S+ or S- events. In order to avoid strong luminance variations at the fixation point during color changes, by default the fixation °point was colored in light grey at a luminance value near to the average luminance of the color changes. We set the duration of the fixation point presentation (ITI = 500 to 750 ms), the first noise placeholder (600 to 1500 ms) and the response time (1500 ms). Participants performed a total of 648 trials divided in 6 blocks of ∼10 minutes. There were a total of 6 experimental conditions of interest: stimulus type (S- | S+ | S++) × modality (“visual” / “audiovisual”), and at the end of the experiment each participant underwent a total of 108 trials per combination of conditions. The contrast level used for the gratings used in the high contrast S++ condition was generated by doubling the contrast estimated for the S+ condition. This condition served to ensure above chance-level decoding classification of signal-present vs signal-absent conditions and to test whether the audiovisual interaction strength changed as a function of the stimulus bottom-up saliency. The vertical grating stimuli where only presented at one eccentricity (center). Since it was impossible to assess whether the subjective visibility of the gratings changed across blocks, as participants did not make decisions about the gratings absence or presence, the contrast value was kept fixed during the whole experiment.

### Behavioral data analyses

In the first experiment, visual sensitivity (d’) and criterion (c) parameters were calculated for each condition and block, applying the signal detection theory to a classic yes-no paradigm (Macmillan & Creelman, 2004). S+ trials correctly reported as “signal” were coded as hits and S- trials incorrectly reported as “signal” were coded as false alarms. In the second experiment the participants’ task was to remember a sequence of items and report whether the third item was the same or different from the previously presented items. Therefore, we applied signal detection theory to calculate the participants sensitivity, but assuming a same-different “independent observation” model paradigm.

### MEG Recording and Preprocessing

Whole-brain neural recordings were registered using a 275-channel MEG system with axial gradiometers (CTF MEG Systems, VSM MedTech Ltd.) located in a magnetically shielded room. Participants’ eye-movements and blinks were tracked online using an EyeLink 1000 (SR Research). Throughout the experiment, head position was monitored online and corrected if necessary using three fiducial coils that were placed on the nasion and on earplugs in both ears. If subjects moved their head more than 5 mm from the starting position, they were repositioned during block breaks. All signals were sampled at a rate of 1,200 Hz. Data preprocessing was carried out offline using FieldTrip (www.fieldtriptoolbox.org). The data were epoched from 2000 ms before and 1500 ms after the signal present / absent event onset. To identify artifacts, the variance (collapsed over channels and time) was calculated for each trial. Those trials and channels with large variances were subsequently selected for manual inspection and removed if they contained excessive and irregular artifacts. Additionally, trials without participant’s response or containing eye-blinks within the interval of 100 ms before or after the target presentation were removed from subsequent analyses. We used independent component analysis to remove regular artifacts, such as heartbeats and eye blinks. For each subject, the independent components were inspected manually before removal. “Bad” channels showing SQUID jumps or other artifacts were interpolated to the weighted by neighbours distance average of neighboring channels. For the main analyses, data were low-pass filtered using a Butterworth filter with a frequency cutoff of 30 Hz and subsequently downsampled to 100 Hz. No detrending was applied for any analysis. Finally, the data were baseline corrected on the interval of −200 ms to the [S+ | S-] event onset (0 ms).

### Source reconstruction and ROI generation

For each participant we build volume conduction models based on single-shell model of the standard Montreal Neurological Institute (MNI) anatomical atlas (Nolte, 2003). Then, we used them to construct search grids (10-mm resolution). For each grid point, lead fields were computed with a reduced rank, which removes the sensitivity to the direction perpendicular to the surface of the volume conduction model.

In order to gain insight on how the auditory stimuli modulate the processing of visual stimuli across the perceptual hierarchy, we generated four neuroanatomically defined regions of interest (ROIs; Fig. 4A) using the AAL anatomical atlas: These were the visual cortex, that is involved in the encoding of sensory information (Ress & Heeger, 2003), the parietal cortex, that is related to evidence accumulation during perceptual decision (Kiani & Shadlen, 2009; Zhou & Freedman, 2019), the inferotemporal cortex, that has been associated to visual memory and target identification (Miller et al., 1993; Mishkin, 1982; Pagan et al., 2013) and the dorsolateral prefrontal cortex, that is involved with short-memory, decision-making and awareness (Funahashi, 2006; Kim & Shadlen, 1999; Philiastides et al., 2011; van Vugt et al., 2018). These brain regions have been typically associated to perceptual decisions in visual detection tasks (see Table 1 in SM). Using the covariance matrix computed from the combined visual and audiovisual trials (-0.2 to 1.5 s, time-locked to the target onset; 10% normalization), the volume conduction model, and the lead field, we applied a linearly constrained minimum variance (van Veen et al., 1997) beamformer approach in Fieldtrip (Oostenveld et al., 2011) to build a common spatial filter for each grid point and participant. Finally, to spatially constrain the analyses to the previously selected neuroanatomical regions, we projected the sensor level ERFs time series through those virtual channels that spatially overlapped with our ROIs.

### Decoding analysis

We applied a multivariate pattern analysis approach to classify single trials as S+ or S- as a function of the neural activity measured from the virtual channels composing each ROI in each time point. The method that we applied was largely based on linear discriminant analysis (Blankertz et al., 2011; Mostert et al., 2015). First, in order to minimize time-point by time-point absolute univariate differences between the visual and audiovisual conditions, we z-scored the ERF activity across virtual channels for each time point and considering the visual and audiovisual conditions independently. Then, using the patterns of activity measured at the multiple virtual channels (i.e. generally termed as features) contained in one of our ROIs, we calculated the weights vector w that optimally discriminates between S+ and S- trials (equation 1).

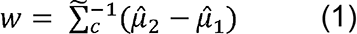

*μ̂*_1_ and *μ̂*_2_ are two column vectors of length F (i.e. number of features) for a given time point, representing the neural activity averaged across the S+ (*μ̂*_2_) and S- (*μ̂*_1_) trials respectively, and 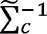 is the common regularized covariance matrix.

The regularization parameter was optimized in preliminary tests using cross-validation and was kept fixed for all subsequent analyses. To make the encoding weights comparable across time points, we added a normalization factor (denominator in equation 2) to the weights vector such that the mean difference in the decoded signal between classes equals a value of one.

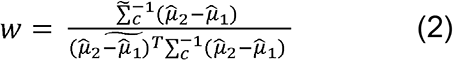

Next, to assess whether the learned weights could discriminate between S+ and S- trials we cross-validated our decoder (equation 2) in a different set of trials X.

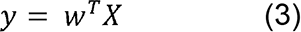

X is a matrix of size F x N, where N represents the number of trials and F the number of features present in the independent dataset. By multiplying X by the weights transpose matrix (N^.l.^ ) that we previously estimated from the training dataset (equation 2), we obtained the decoder output, termed here as the “discriminant channel”. If there is information in the neural signal pertaining to the classes to be decoded, we expect the mean discriminant channel amplitude to be *ȳ*_2_ > *ȳ*_1_, whereas if no information is available in the neural signals, we must find *ȳ*_1_ = *ȳ*_2_. In order to classify a trial as S+ or S-, we set a cut-off value. If the difference between the discriminant channels *ȳ*_2_ and *ȳ*_1_ was larger than 0, the trial was classified as S+. Instead, if the difference between the discriminant channels *ȳ*_2_ and *ȳ*_1_ was smaller than 0, the trial was classified as S-. To avoid “double dipping” (Kriegeskorte et al., 2009) we adopted a leave-one-out cross-validation approach where we randomly divided the trials of the dataset in five evenly distributed manifolds. We built an unbiased classifier by training our discriminant model in four of the five manifolds that contained visual and audiovisual trials. Subsequently, we tested the classifier in the remaining manifold. We repeated the same process until all the manifolds were used as training and test sets. Moreover, given potentially unequal trial numbers for each visual and audiovisual condition, we repeated the same process 50-times and averaged the final discriminant channel output for each trial.

We calculated sensitivity and criterion parameters from neural activity. For instance, if the classifier categorized a S+ trial as S+, we coded the trial as a hit. Instead, if the classifier wrongly categorized a S- trial as S+, we coded the trial as a false alarm. Using the same approach as in a signal detection theory Yes-No paradigm (Green & Swets, 1966), we estimated sensitivity (d’) and criterion (c) parameters for each time point in each trial based on neural activity derived hits and false alarms. The d’ parameter allowed us to evaluate the sensitivity of the classifier to discriminate between S+ and S- trials in the visual and audiovisual conditions. Time points with d’ > 0 represents above chance level visual stimuli classification. On the other hand, the c parameter indexed the bias of the classifier to classify the trials as S+ or S- in the visual and audiovisual conditions. Specifically, time points with lower c values represent stronger biases to report the signal presence regardless the actual presence or absence of the signal. This procedure allowed us to directly compare participants behavioral performance based on signal detection theory parameters with the same parameters estimated from neural activity during the perceptual decision making.

The decoding analysis outlined above was implemented in a time-resolved manner by applying it sequentially at each time point in steps of 10 ms, resulting in a decoders array of the same length as the number of trial time points. To characterize the temporal organization of the neural processes that underlie the auditory contributions to visual processing (Fig. 2), we implemented a temporal generalization (TG) method (King & Dehaene, 2014). Each decoder trained on any specific time point was applied to all other time points. If we average the decoder output over trials, this results in a squared temporal generalization matrix (TGM) with “training time” × “decoding time” values per condition. The diagonal values in the TGM contain the estimated parameters for the decoders trained and cross-validated in the same time points (t_train_ = t_test_). Instead, the row values in the TGM represent how a specific decoder trained at a time point t_train_ in both visual and audiovisual conditions classifies visual and audiovisual trials as signal-present or signal-absent trials at earlier and later time points. In addition, a column gives insight into whether the neural patterns of activation of the two conditions can are discriminated at time point t_test_ on the basis of the decoders trained on all other time points. In summary, observing that the same decoder can separate between conditions at multiple time points give us relevant information about how persistent in time neural representations are.

### Statistical analyses

#### Cluster-based permutation tests

To statistically assess in which training and testing time combinations of the TGM the decoder successfully discriminated the trials as S+ or S- above chance level, we contrasted the d’ values against 0. We applied cluster-based permutation tests (Maris & Oostenveld, 2007). This procedure controls for multiple comparisons across decoding and testing time combinations by leveraging the inherently correlated nature of neighbouring observations. For each TGM condition, each d’ value was compared univariately against 0 using a t-score. Positive and negative clusters were then formed separately by grouping temporally adjacent data points whose corresponding P-values were lower than 0.05 (two-tailed). Cluster-level statistics were calculated by summing the t-values within a cluster, and a permutation distribution of this cluster-level test statistic (1000 permutations per contrast) was computed. The null hypothesis was rejected for those clusters for which the p-value were smaller than 0.05, compared to the permutation distribution. In order to compare how sounds modulated d’ and c at different time points, we also applied cluster-based permutation tests but contrasting the visual and audiovisual TGMs with each other.

## Supporting information

Supplemental analyses

## Acknowledgments

We are very grateful to Pim Mostert and Valentin Wyart for sharing their code and initial advice on the MEG and behavioral analyses. We thank Salvador Soto-Faraco for helpful comments on an earlier draft of the manuscript. This work was supported by the Netherlands Organisation for Scientific Research (NWO Veni grant 016.Veni.198.065 awarded to ES and Vidi grant 452-13-016 awarded to FPdL), the EC Horizon 2020 Program (ERC starting grant 678286 awarded to FPdL) and the AGAUR Generalitat de Catalunya (Beatriu de Pinòs 2017-BP-00213 awarded to APB). Author contributions: A.P.B, and F.L. conception and design of research; A.P.B. performed experiments; A.P.B and E.S. analyzed data; A.P.B. drafted manuscript; A.P.B., E.S., and F.L. edited, revised and approved the final version of the manuscript.

