## Supplemental analyses for "Seeing sounds: Neural mechanisms underlying auditory contributions to visual detection"

Supplemental Information Inventory:

1. Methods: ROIs definition
   1. Fig. S1
   2. Table 1
2. Experiment 1: Participants hits and false alarms
   1. Fig. S2
3. Experiment 2: Participant’s performance
   1. Fig. S3
4. Decoder hits and false alarms in experiment 1 & 2
   1. Fig. S4
   2. Fig. S5
5. Experiment 1: Neural decoder’s sensivity and criterion parameters and their correlation with behavioral performance.
   1. Fig. S5
   2. Fig. S6
6. Experiment 1: Control analyses
   1. Fig. S7
   2. Fig. S8
7. Experiment 2: Neural decoder sensitivity and criterion parameters in Low contrast condition
   1. Fig. S9
   2. Fig. S10
8. Experiment 2: Univariate modulations in the visual cortex ROI
   1. Fig. S11
9. Experiment 1: Temporal ordering of clusters across ROIs
   1. Fig. S12

**Methods: ROIs definition**

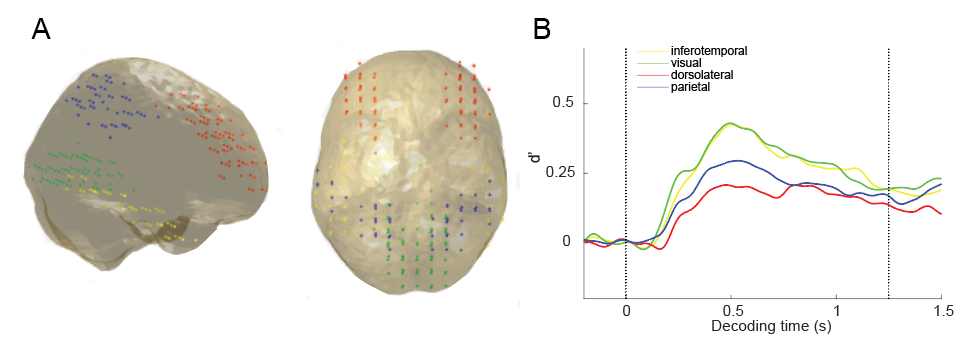

Figure S1.A Grid points conforming each ROI overlayed on two different views of a MNI standard template headmodel (Visual cortex ROI: green points, Inferotemporal cortex: yellow points, Parietal cortex: blue points, Dorsolateral prefrontal cortex: red points). B. Decoding sensitivity in Experiment 1 (visual and audiovisual modalities together) for each ROI at matching training and decoding time points (diagonal of the TGM): Information decoding peaks at 500 ms and nicely follows the temporal ordering expected for each ROI as a function of its level within the perceptual hierarchy.

Table 1

|  | N | AAL label | AAL label |
| --- | --- | --- | --- |
| Visual cortex | 67 | Calcarine L and R | Lingual L and R |
| Parietal cortex | 53 | Parietal Superior L and R | Parietal Inferior L and R |
| Interotemporal cortex | 52 | Temporal Inferior L and R |  |
| Dorsolateral prefrontal cortex | 79 | Frontal Middle L and R |  |

Table listing the number of gridpoints and AAL atlas labels conforming each anatomically defined region of interest.

**Experiment 1: Participants hits and false alarms**

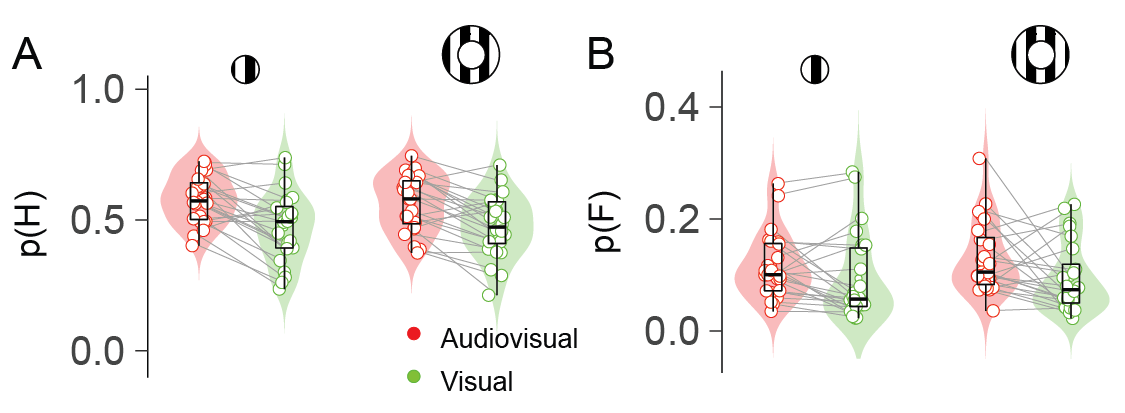

Figure S2. Behavioral results in experiment 1: Proportions of hits (A) and false alarms (B) are depicted for each modality (“audiovisual”, “visual”), visual eccentricity condition (“center” and “periphery”) and participant. Inside the violin plot, the horizontal black line reflects the median, thick box indicates quartiles, and whiskers 2 ´ the interquartile range. Grey horizontal lines connect participants results across conditions.

**Experiment 2: Participant’s performance**

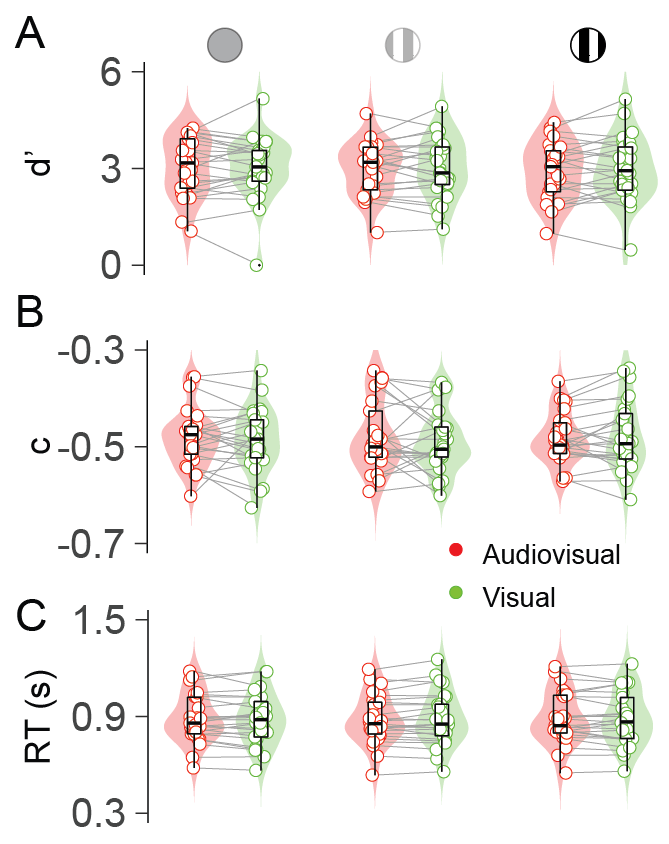

Figure S3. Behavioral results in the memory task of experiment 2: Sensitivity (A), criterion (B) and reaction times (C) are represented for each modality (“audiovisual”, “visual”), distractor contrast (“S-”, “S+” and “S++” signal) and participant. Same labelling convention as in Figure S.

**Decoded hits and false alarms**

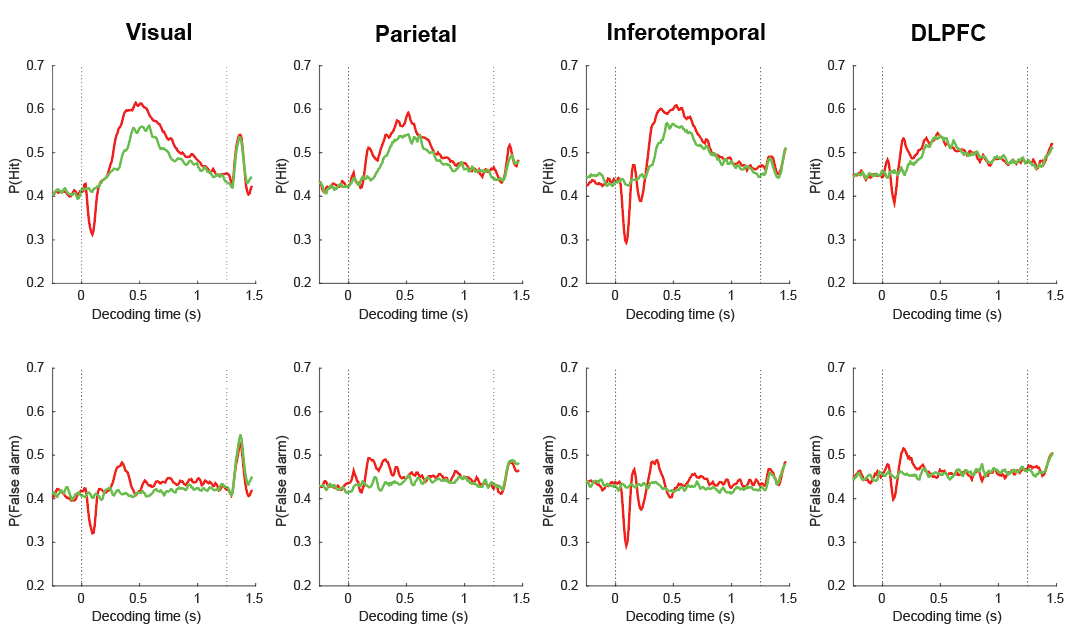

Figure S4. Proportion of hits (upper panels) and false alarms (bottom panels) estimated by a classifier trained at the decoding peak (500 ms) in experiment 1. The visual and audiovisual modalities are represented with green and red lines respectively.

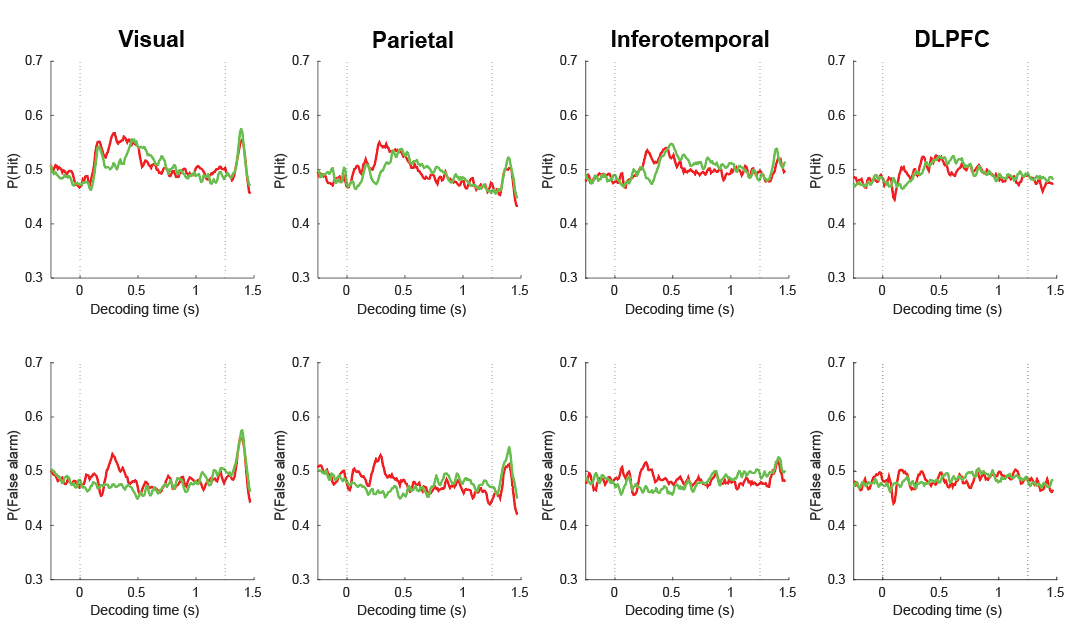

Figure S5. Proportion of hits (upper panels) and false alarms (bottom panels) estimated by a classifier trained at (500 ms) in experiment 2. The visual and audiovisual modalities are represented using green and red lines respectively. See that there is a larger proportion of decoded false alarms at 200 to 350 ms in the audiovisual compared to the visual condition.

**Experiment 1: Neurally decoded sensitivity and criterion parameters and their correlation with behavioral performance**

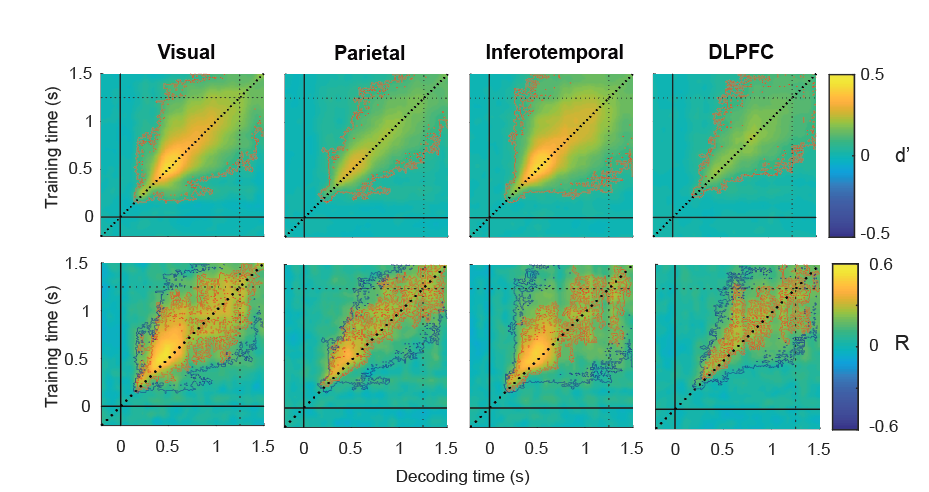

Figure S5. Upper panels: Decoded sensitivity parameters in each ROI (considering the visual and audiovisual conditions together). Bottom panels: Time point by time point correlation between the six neurally and behaviorally estimated d’. Red thin contours depict significant clusters. Blue thin contours (in the bottom panel) overlay the significant sensitivity decoding clusters in the upper panels. Most of significantly decoded information delimited by the blue contour was correlated with participants performance.

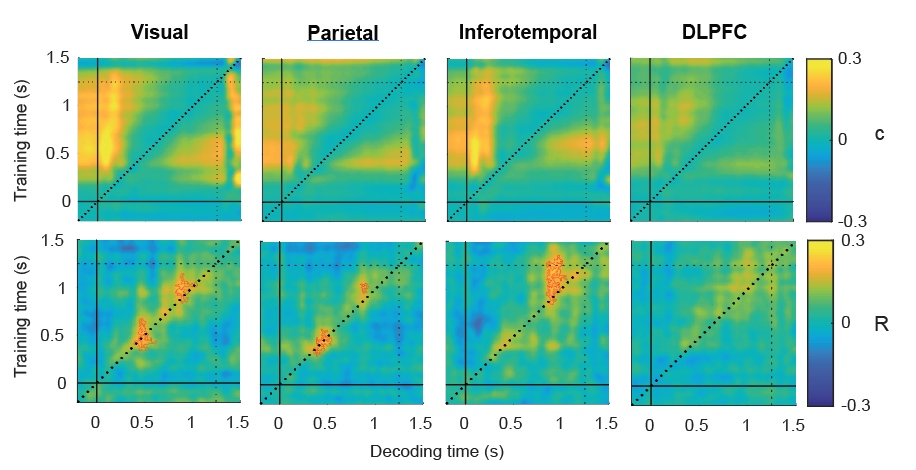

Figure S6. Upper panels: Decoded criterion parameters in each ROI (considering the visual and audiovisual conditions together). Bottom panels: Time point by time point correlation between the six neurally and behaviorally estimated c parameters. Red thin contours depict significant clusters.

**Experiment 1: Control analyses**

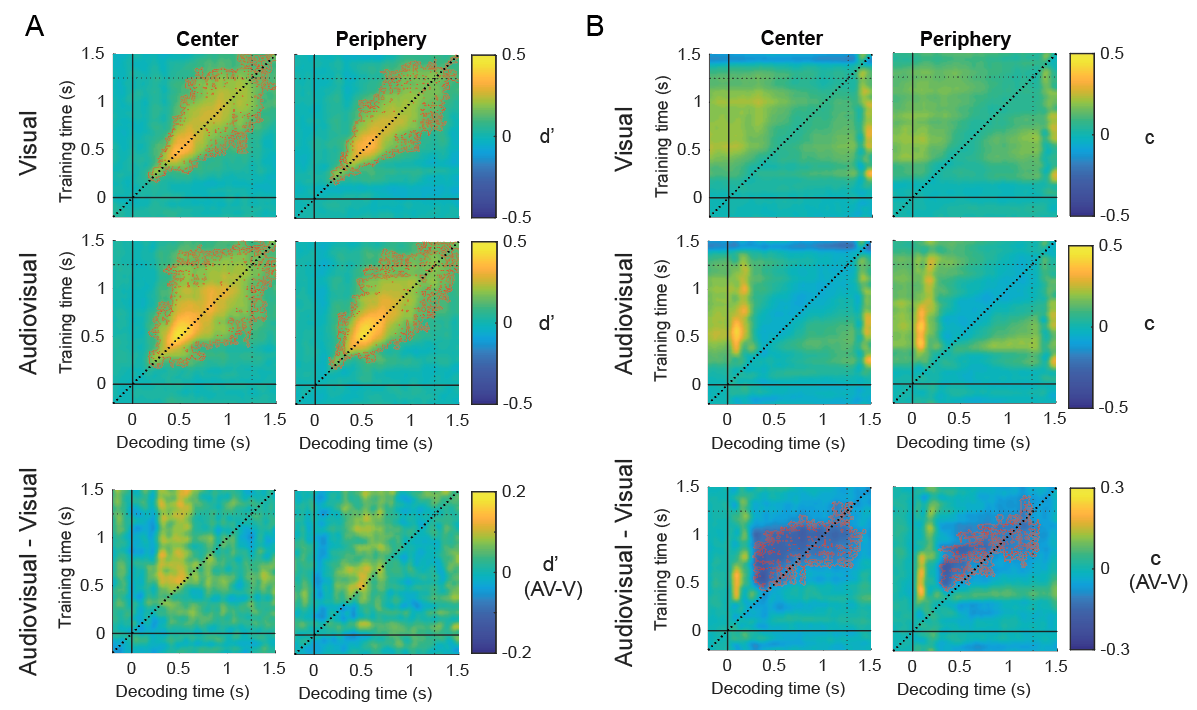

Figure S7. Temporal generalization matrixes depict (A) the neurally decoded sensitivity parameters (d’) in the visual (upper TGMs) and audiovisual conditions and their difference, after training and testing the classifiers with data from the center (left panels) and periphery (right panels) visual field conditions (only visual cortex ROI). We found qualitatively similar patterns of enhanced decoding at center and periphery visual fields. We did not find significant clusters, probably due the small number of trials used to train and test the classifiers but in essence these results demonstrate that the vertical enhanced generalization pattern is present at center and periphery visual fields. B. illustrates similar results as A but using criterion parameters.

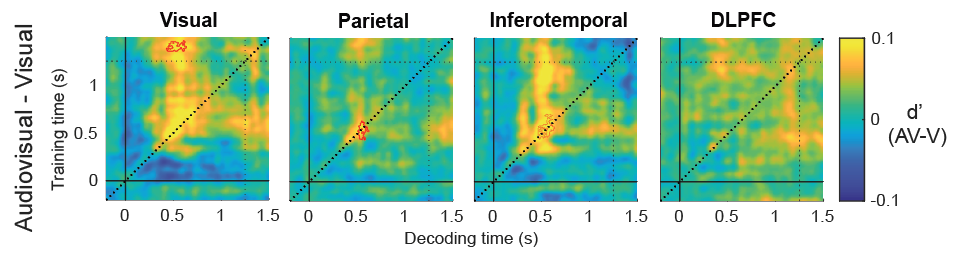

Figure S8. Differential d’ temporal generalization matrixes generated by training and testing the decoders in the visual and audiovisual conditions separately. This control analysis recovers the expected symmetry with respect to the diagonal (unity line), that we did not observe when the decoders were trained in both modalities together.

**Experiment 2: Decoded sensitivity and criterion parameters in Low contrast condition**

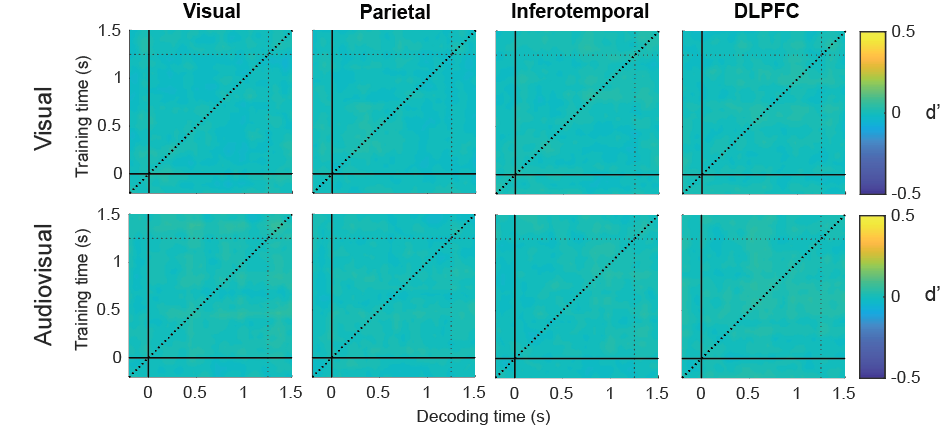

Figure S9. Temporal generalization matrixes depict (A) the neurally decoded sensitivity parameters (d’) in the low contrast visual (upper TGMs) and audiovisual (bottom TGMs) conditions for each ROI (Experiment 2).

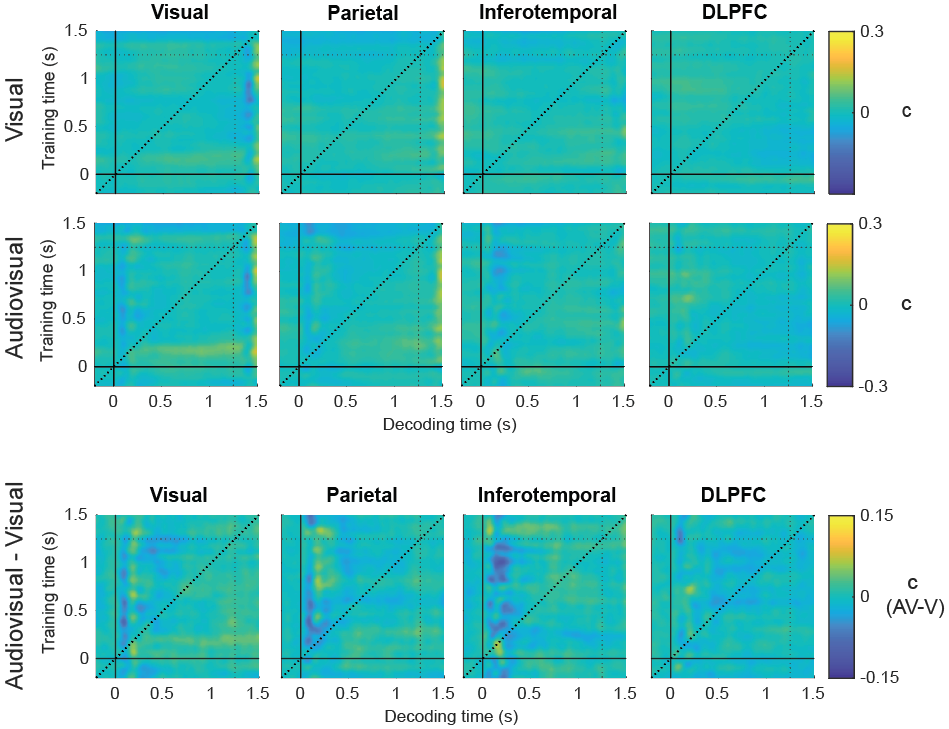

Figure S10. Temporal generalization matrixes depict (A) the neurally decoded criterion parameter (c) in the low contrast visual (upper TGMs) and audiovisual (bottom TGMs) conditions, and their difference for each ROI (Experiment 2).

**Experiment 2: Univariate modulations in the visual cortex ROI**

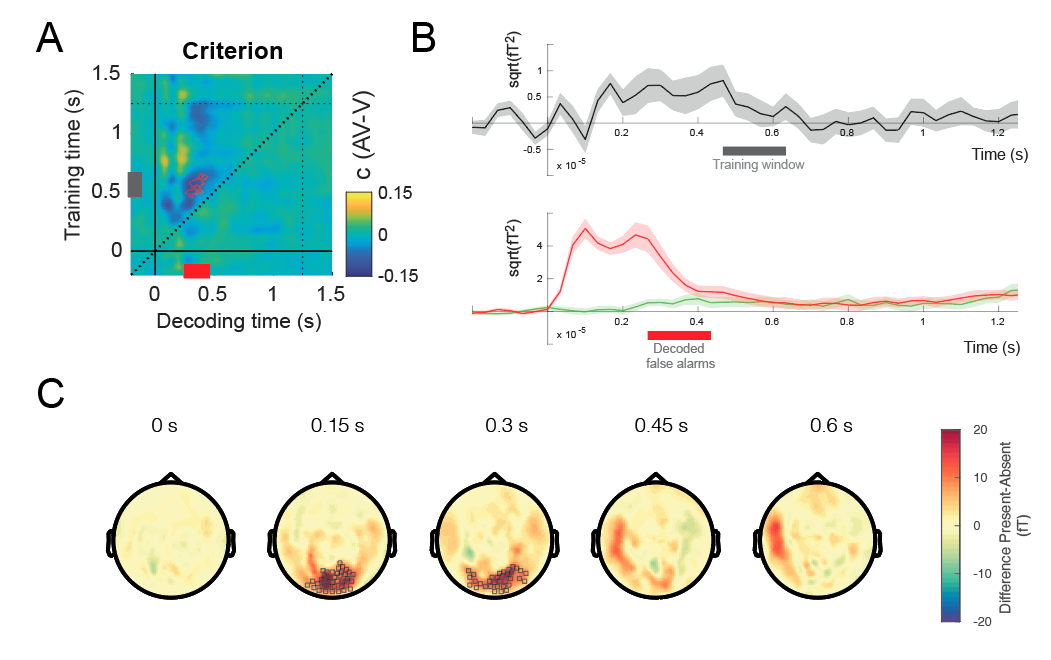

Figure S11. Stimulus-induced univariate modulations as a function of differences in neurally decoded criterion in the visual cortex ROI. A. Temporal generalization matrix showing the cluster containing the significant difference in criterion between the audiovisual and visual conditions in the visual cortex ROI. Grey and red rectangles depict the temporal windows in which multiple classifiers were trained and tested, leading to a systematic positive bias in signal presence classification (i.e. criterion reduction). B represents the difference between S++ and S- source reconstructed ERF activity in the visual ROI (averaged over virtual channels) regardless of modality (upper panel), and the ERF activity measured in the audiovisual (red) and visual (green) conditions (bottom panel). C Group-level scalp topography for the high contrast condition showing sensor-level S++ against S- ERF differences (averaging over visual and audiovisual conditions). Black boxes represent those MEG sensors that conformed a significant cluster in each particular time point.

**Experiment 1: Temporal ordering of clusters across ROIs**

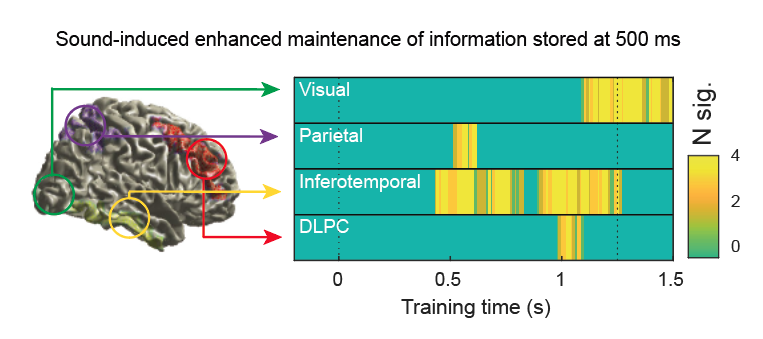

Figure S12. The information encoded at 500 ms can be better decoded in the audiovisual compared to the visual condition at different latencies in each ROI. Here we illustrate the temporal ordering of the significant cluster across the different ROIs. Although these results do not allow to make strong claims about the spatial dynamics of the sound-induced visual enhancement, they can inspire new testable hypotheses. Colorbar represents the total number of significant timepoints within a temporal window of 0.45 to 0.55 s in the “decoding time” axis along the “training time” axis in Figure 4. The two vertical dotted lines represent the stimulus onset and the beginning of the response phases respectively.
